# Lactate potentiates differentiation and expansion of cytotoxic T cells

**DOI:** 10.1101/571745

**Authors:** Helene Rundqvist, Pedro Veliça, Laura Barbieri, Paulo A. Gameiro, Pedro P. Cunha, Milos Gojkovic, Sara Mijwel, Emil Ahlstedt, Iosifina P. Foskolou, María Victoria Ruiz-Pérez, Marie Arsenian-Henriksson, Jernej Ule, Arne Östman, Randall S. Johnson

## Abstract

Exercise has a range of effects on metabolism. In animal models, repeated exertion reduces malignant tumour progression, and clinically, exercise can improve outcome for cancer patients. The etiology of the effect of exercise on tumour progression is unclear, as are the cellular actors involved. We show here that exercise-induced reduction in tumour growth is dependent on CD8+ T cells and that lactate, which is produced at high levels during exertion, increases proliferative capacity and cytotoxicity of CD8+ T cells. We found that at elevated levels lactate is used as a fuel during T cell activation. We further found that injection of lactate into animals can reduce malignant tumour growth in a dose-and CD8+ T cell-dependent manner. These data demonstrate that lactate can act to increase the anti-tumour activity of cytotoxic T cells, and in so doing, reduce cancer progression.

## Introduction

In humans, exercising cohorts have lower rates of cancer incidence^1^, and better outcomes across a range of cancer diagnoses^2,3^, proportionate to the degree and intensity of exercise. The mechanisms underlying these observations have remained elusive, although recent work has indicated a relationship between immune response and exercise-induced changes in malignant progression indicating that this could be due to NK cell function^4^, or innate immune cells more generally^5^.

The metabolic demands of physical exertion generally induce an enhanced and significantly glycolysis-derived energy production. The metabolic perturbation caused by exertion can cause a change in the ratio of energy substrates utilized and shift metabolite profiles in ways that act to support energy production throughout the body^6^.

Within the adaptive immune system, T cells play a crucial role in controlling tumour growth. By recognising mutation-derived neoantigens, T cells identify and eliminate malignant cells in a process known as immunosurveillance^7,8^. Escape from immune control is a critical step toward progressive malignant growth in many cancers, and tumours achieve this in a number of ways, including the dampening of anti-tumour T cell responses through immunosuppressive mechanisms such as expression of CTLA-4 and PD-L1.^7–10^.

Thus, an inhibited endogenous anti-tumour T cell response exists within many patients with progressive disease. Some cancer immunotherapies, e.g., checkpoint blockade therapy with monoclonal antibodies against CTLA-4 and PD-1/PD-L1, can create an anti-tumour T cell response^11^. In others, anti-tumour T cells are generated *ex vivo* by expanding tumour-infiltrating lymphocyte (TIL) populations or using gene transfer to express defined T cell receptors (TCR) or chimeric antigen receptors (CAR) in patient-derived T cells, and adoptively transferring them back into the patient^12-15^. Improving the fitness, expansion and function of T cells is central to a favourable prognosis and the success of cancer immunotherapy.

The activity of immune cells is tightly linked with their metabolism, both at an intrinsic and extrinsic level^16,17^. In this study we investigate the association between exercise-derived metabolites and CD8+ T-cell function. To address this, we undertook studies of exercise-induced changes in tumour progression, and asked whether lactate, the most substantially increased metabolite from exercising muscle, has a role in CD8+ T cell function. We show that lactate is taken up by CD8+ T cells and used as a carbon fuel to drive both effector differentiation and proliferation and show that this has clear immunotherapeutic implications.

## Results

### Depletion of CD8+ T cells abolishes the anti-tumoural effects of exercise

To address the role of immunity in the effects of exercise on neoplasia, we first assessed the effect of repeated voluntary exertion on tumour progression in a genetic mouse model of breast cancer induced by the MMTV-PyMT transgene, placed on the FVB inbred strain background. FVB inbred mice are enthusiastic runners relative to other inbred strains^18^ and the MMTV-PyMT model mimics in many regards the gradual progression of human breast cancer^19^. Exercising MMTV-PyMT transgenic mice showed an onset of palpable tumours at an average age of 49 days, which was comparable to their sedentary counterparts, whose onset was at an average age of 50 days (Supplementary Figure 1A). Contrary to what was previously shown^20^, the running mice showed no statistically significant differences in tumour growth between the groups (Supplementary Figure 1B, C and D). However, the infiltration of Granzyme B (GzmB) positive cells was significantly higher in primary tumours of running mice, while voluntary running had no effect on the frequency of CD3, F4/80, PCNA or podocalyxin-expressing cells (Supplementary Figure 1E), indicating that exercise in this tumour model specifically induced the infiltration of cytotoxic lymphocytes and not the infiltration of stromal cell types generally.

Given that infiltration of cytotoxic T cells and NK cells is linked to a favourable prognosis in many human neoplasms and in some animal tumour models^21^, we proceeded to explore the role of lymphoid cells, and in particular cytotoxic CD8+ T cells, in the process of exercise-induced modulation of tumour growth. As the transgenic MMTV-PyMT model may be predominantly affected by myeloid cells^19^, in the next set of experiments, immunocompetent FVB mice were given access to an exercise wheel 14 days prior to, and then following, subcutaneous injection with a murine mammary cancer line derived from the MMTV-PyMT model, the I3TC cell line, which may trigger a different stromal response from that seen in the transgenic model^22^. In this cell line-driven model, allowing animals to exercise significantly reduces tumour progression and increases survival times (Figure 1A) relative to animals not allowed access to an exercise wheel.

**Figure 1.**
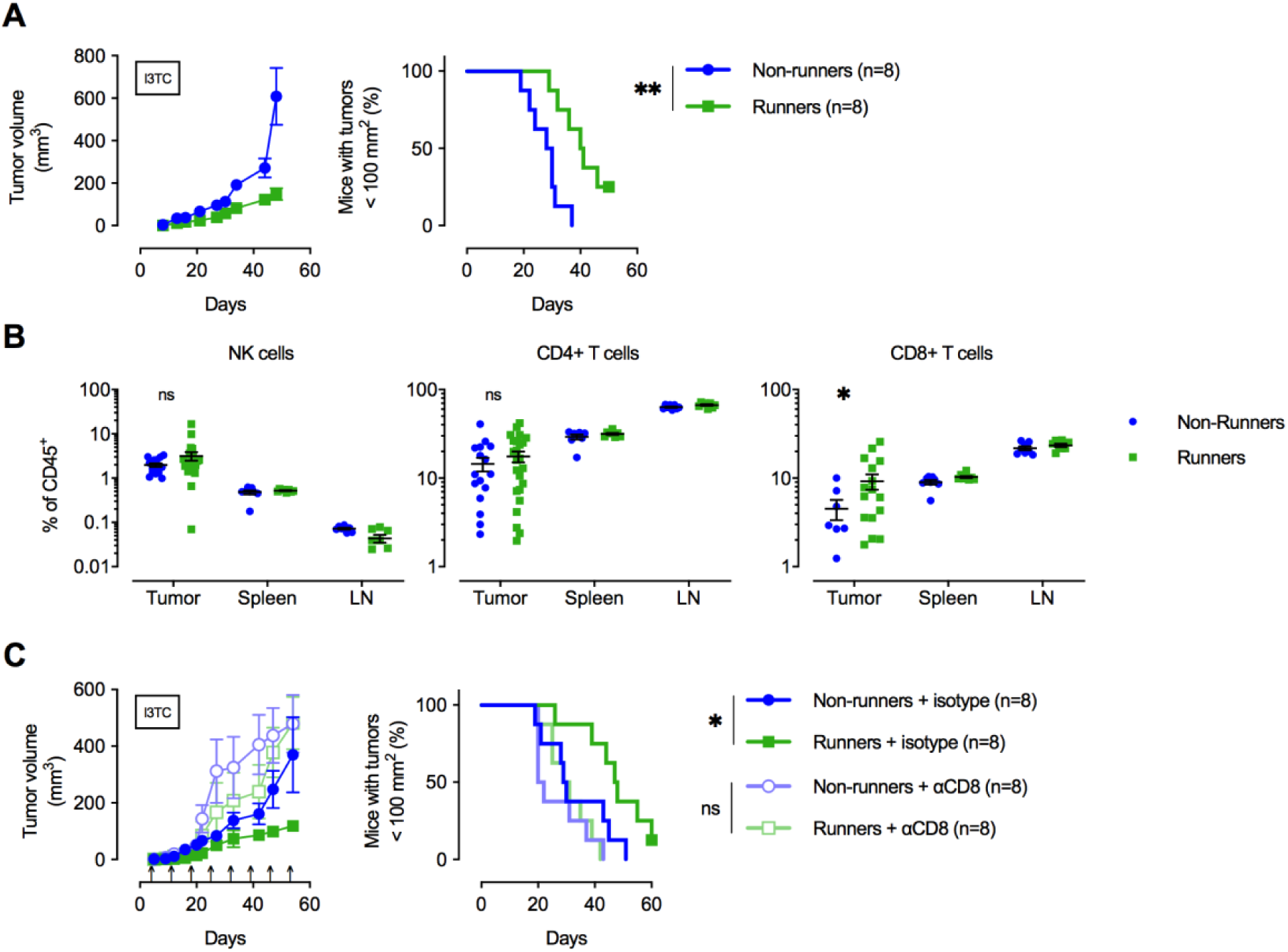
Exercise delays tumour growth in a CD8+ T-cell dependent manner. **A.** FVB mice were allowed to exercise voluntarily (in running wheels, runners) or left non-exercised (locked running wheels, non-runners) before and after being inoculated subcutaneously with 5×10^5^ tumour cells of the breast cancer cell line I3TC. Graphs show mean tumour volume and SEM over time (left) and survival (right). ** P <0.01, Log-rank (Mantel-Cox) survival test. **B.** Flow cytometry determined frequency of lymphocytic populations within I3TC tumour, spleen and draining lymph node (LN) at day 55 after inoculation. * P <0.05 column factor in a two-way ANOVA. ns= not significant. **C.** Same experimental setting as in (A) with or without weekly antibody-mediated depletion of CD8+ T cells (arrows). Graphs show mean tumour volume and SEM over time (left) and survival (right). ** P <0.01, * P <0.05, Log-rank (Mantel-Cox) survival test.

To determine whether immune cell populations were also affected by exercise in the subcutaneous I3TC model, we carried out flow cytometric analysis of single cell suspensions of inoculated tumours, as well as spleens and tumour-draining lymph nodes of running versus non-running animals. In this analysis, only CD8+ T cells showed a significant increase in frequency across these three tissues in the running animals (Figure 1B), while no significant changes were seen in the frequency of CD4+ T cells, NK cells (Figure 1B), macrophages or neutrophils (Supplementary Figure 1F). To address the role of increased CD8+ T cell populations in the reduction of tumour growth caused by exercise, we depleted these cells by weekly injections of anti-CD8 antibodies in exercising and non-exercising mice during tumour growth. CD8+ T cell depletion (Supplementary Figure 1G) significantly reduced the beneficial effects of exercise on tumour growth and long-term survival (Figure 1C), demonstrating a role for cytotoxic T cells in the suppression of tumour growth by exercise.

### Lactate alters differentiation and expansion of CD8+ T cells

Exercise causes changes in a wide range of metabolic mediators of immune response^6^. However, the single most profound metabolic change induced by exertion is the transient increase in circulating lactate. Circulating lactate levels increase very rapidly during exercise, and can in response to high intensity exertion rise up to 100-fold in metabolizing tissues such as skeletal muscle^23,24^, with a concomitant increase of more than 10-fold in plasma^25^. As the data above demonstrates, exercise reduces tumour growth in a CD8+ T cell-dependent manner; we thus sought to determine whether lactate, the predominant metabolite generated during exercise, could be a determining factor in the action of CD8+ T cells in this process.

To test this, we activated CD8+ T cells *ex vivo* for 3 days in the presence of increasing doses of lactic acid, sodium lactate, or sodium chloride. We used both lactic acid and sodium lactate to distinguish between the effects of the lactate anion released during glycolysis, and the actual acidification caused by proton dissociation (Supplementary Figure 2A). Although the production of lactic acid is the immediate consequence of increased glycolytic flux during exercise, in healthy tissues, acidification is rapidly buffered in surrounding tissue and in plasma^26^. For this reason, lactate produced during exercise does not alter circulating plasma pH levels in the manner seen locally in, for example, solid tumours^27^. As expected, addition of lactic acid to culture media causes a significant drop in pH, while sodium lactate or sodium chloride do not alter media pH (Supplementary Figure 2B). To better understand the role of sodium lactate, while controlling for osmolality (Supplementary Figure 2C), we employed equimolar amounts of sodium chloride as an osmolality control in subsequent experiments.

As reported previously^27^, pH-lowering lactic acid has deleterious effects on CD8+ T-cell activation, resulting in reduced cell numbers and a disruption of GzmB upregulation (Figure 2A and B). In contrast, activation in the presence of pH-neutral sodium lactate resulted in a dose-dependent increase in GzmB expression, beginning at concentrations of 10 mM (Figure 2A and B). Interestingly, activation in the presence of pyruvate, a metabolite generated by lactate oxidation, caused an impaired CD8+ T cell accumulation and lowered GzmB induction (Supplementary Figure 2D).

**Figure 2.**
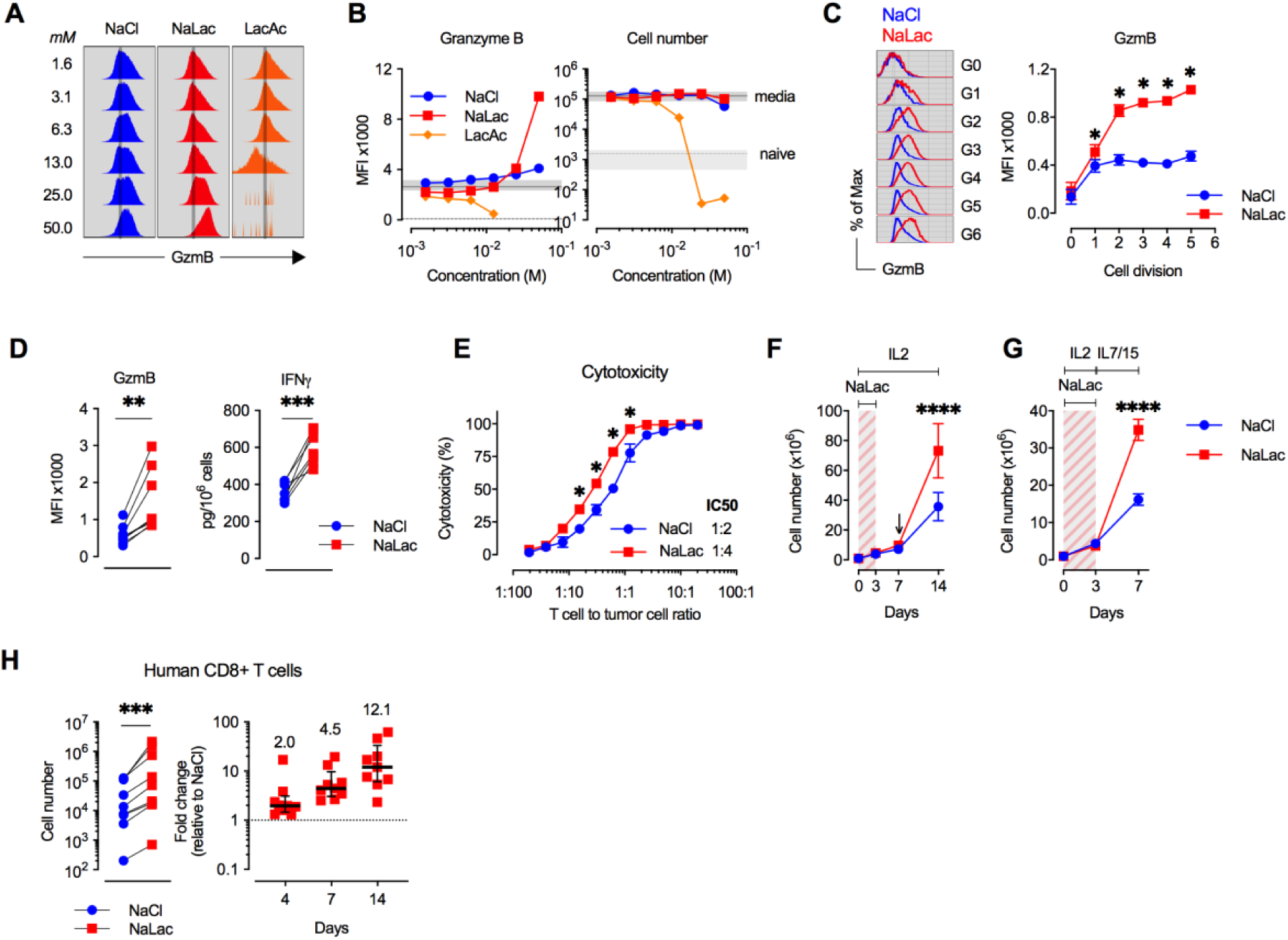
Lactate boosts proliferation and cytotoxic differentiation of CD8+ T cells. **A.** Mouse CD8+ T cells were activated for 72 hours in the presence of varying concentrations of Sodium Chloride (NaCl), Sodium Lactate (NaLac) or Lactic Acid (LacAc). Flow cytometry histograms showing intracellular granzyme B (GzmB) levels. **B.** Median Fluorescence Intensity (MFI) of GzmB and cell number after activation in the presence of NaCl, NaLac and LacAc. Solid line: average of non-treated CD8+ T cells; Dotted line: average of non-activated T cells. Grey area: range (minimum to maximum) of non-treated or non-activated controls. **C.** MFI of GzmB at each cell generation (G0-G6) of mouse CD8+ T cells activated as in (C). Left: Flow cytometry histograms of NaCl (blue) and NaLac (red)-treated cells. Right: Mean and SEM of n=6 independent mouse donors. * P <0.0001 repeated-measures two-way ANOVA with Sidak’s multiple comparison test. **D.** Levels of secreted interferon-γ (IFN-γ) and MFI of GzmB in mouse CD8+ T cells activated for 72h in the presence of 40 mM NaCl or 40 mM NaLac. Each pair represents an independent mouse donor (n=7-9). *** P<0.001, ** P<0.01, two-tailed paired t-test. **E.** Cytotoxicity against EL4-OVA tumour cells by OVA-specific OT-I CD8+ T cells activated for 3 days in the presence of 40 mM NaCl or NaLac. Graph represents specific cytotoxicity of n=3 independent mouse donors at varying effector-to-target ratios. * P <0.01 repeated-measures two-way ANOVA with Sidak’s multiple comparison test. **F.** Cell number of mouse CD8+ T cells activated in the presence of interleukin-2 (IL-2) and 40 mM NaCl or NaLac. At day 3 cells were washed and cultured with IL-2 henceforth. Cells were re-stimulated at day 7 (arrow) and cultured until day 14. n = 3 independent mouse donors. **** P <0.0001 repeated-measures two-way ANOVA with Sidak’s multiple comparison test. **G.** Cell number of mouse CD8+ T cells activated in the presence of IL-2 and 40 mM NaCl or NaLac. At day 3 cells were washed and cultured with IL-7 and IL-15 henceforth. n = 9 independent mouse donors. **** P <0.0001 repeated-measures two-way ANOVA with Sidak’s multiple comparison test. **H.** Human CD8+ T cells were purified from peripheral blood and activated for 4 days with anti-CD3/CD28 beads and IL-2 in the presence of 40 mM NaCl or NaLac. At day 4 cells were washed and cultured henceforth in the presence of IL-2 alone. Cells were counted at days 4, 7 and 14. Left: Cell number at day 14. Each pair represents an independent human donor (n=8). *** P<0.001, two-tailed paired t-test. Right: fold difference between NaLac and NaCl treatment for each donor at days 4, 7 and 14.

Accumulation of intracellular GzmB was augmented by sodium lactate from the first cell division and remained high in following cell generations (Figure 2C). Increased GzmB after exposure to 40 mM sodium lactate was consistently observed in CD8+ T cells from several mouse donors, as was an increase in secreted interferon-γ (IFNγ), crucial effector molecules in cytotoxic T cell function (Figure 2D). Sodium lactate also caused a small increase in cell size, and a decrease in total CD8+ T cell number due to a slight reduction in proliferation (Supplementary Figure 2E and G), after three days of exposure to the metabolite. Expression of a number of other proteins involved in CD8+ T cell differentiation was modulated after 72 hours in 40 mM sodium lactate: these changes included increased levels of CTLA-4, and 4-1BB, and decreased levels of T-bet, CD62L, CD27 and PD-1 (Supplementary Figure 2F, H, I, and L).

GzmB is the most highly expressed effector protein in activated mouse CD8+ T cells^28^ and is essential to granule-mediated apoptosis of target cells^29^. In keeping with this, CD8+ T cells activated in the presence of sodium lactate for 72 hours showed increased cytotoxicity against tumour target cells (Figure 2E, Supplementary Figure 2N).

Continuous exposure to sodium lactate after day 3 did not impair CD8+ T cell proliferation (Supplementary Figure 2J), but decreased GzmB and increased CD62L expression relative to controls after 7 days of exposure (Supplementary Figure 2K). However, if sodium lactate was withdrawn after day 3, effector protein expression remained largely unchanged by day 7 (Supplementary Figure 2L and M). In addition, limiting sodium lactate exposure to the first 3 days of activation resulted in a 2-fold enhancement of CD8+ T cell expansion in the presence of IL-2 by day 14 (Figure 2F and Supplementary Figure 2J), and a 3-fold enhancement of expansion in the presence of IL-7 and IL-15 by day 7 (Figure 2G).

To determine if these effects on differentiation and proliferation were also seen in human CD8+ T cells, we purified naïve CD8+ T cells from peripheral blood of 9 healthy donors, where lactate exposure was limited to the first 4 days of activation (Figure 2H and Supplementary Figure 2O and P). By day 14, lactate-exposed CD8+ T cells had expanded 12-fold compared with cells from the same donors treated with sodium chloride (Figure 2H). However, unlike what was observed in murine CD8+ T cells, lactate did not alter expression of differentiation markers at this stage, including GzmB (Supplementary Figure 2P). The remarkable proliferative boost during CD8+ T cell *ex vivo* culture not only uncovers a potential role for lactate in driving T cell growth, but also has implications for modulating the production of therapeutic T cells.

Generation of therapeutic T cells for adoptive cell transfer into cancer patients requires extensive *ex vivo* culturing and expansion of patient-derived T cells^13–15^. Thus, we employed a murine model of T cell engineering to test if lactate can increase the production of therapeutic T cells. Mouse polyclonal CD8+ T cells were activated *ex vivo* in the presence of 40 mM sodium lactate or sodium chloride for 3 days, at which point retroviral gene transfer was employed to introduce the alpha and beta chains of the ovalbumin (OVA)-specific OT-I TCR (Figure 3A, Supplementary Figure 3A). Cells were cultured for 4 more days after transduction without lactate or sodium chloride. By day 7, the proportion of cells expressing the OT-I TCR at the surface was higher by 1.5-fold in lactate-treated T cells, while the absolute number of TCR-transduced CD8+ T cells was 3-fold higher than controls (Figure 3B, C and Supplementary Figure 3B). We then employed a mouse model of adoptive cell transfer and injected equal numbers of OT-I-transduced CD8+ T cells into mice bearing subcutaneous ovalbumin-expressing Lewis Lung Carcinoma (LLC-OVA, Figure 3D). Lactate-treated OT-I-transduced cells were as potent as control-treated cells in delaying tumour growth (Figure 3E) and in peripheral blood expansion (Figure 3F and Supplementary Figure 3C).

**Figure 3.**
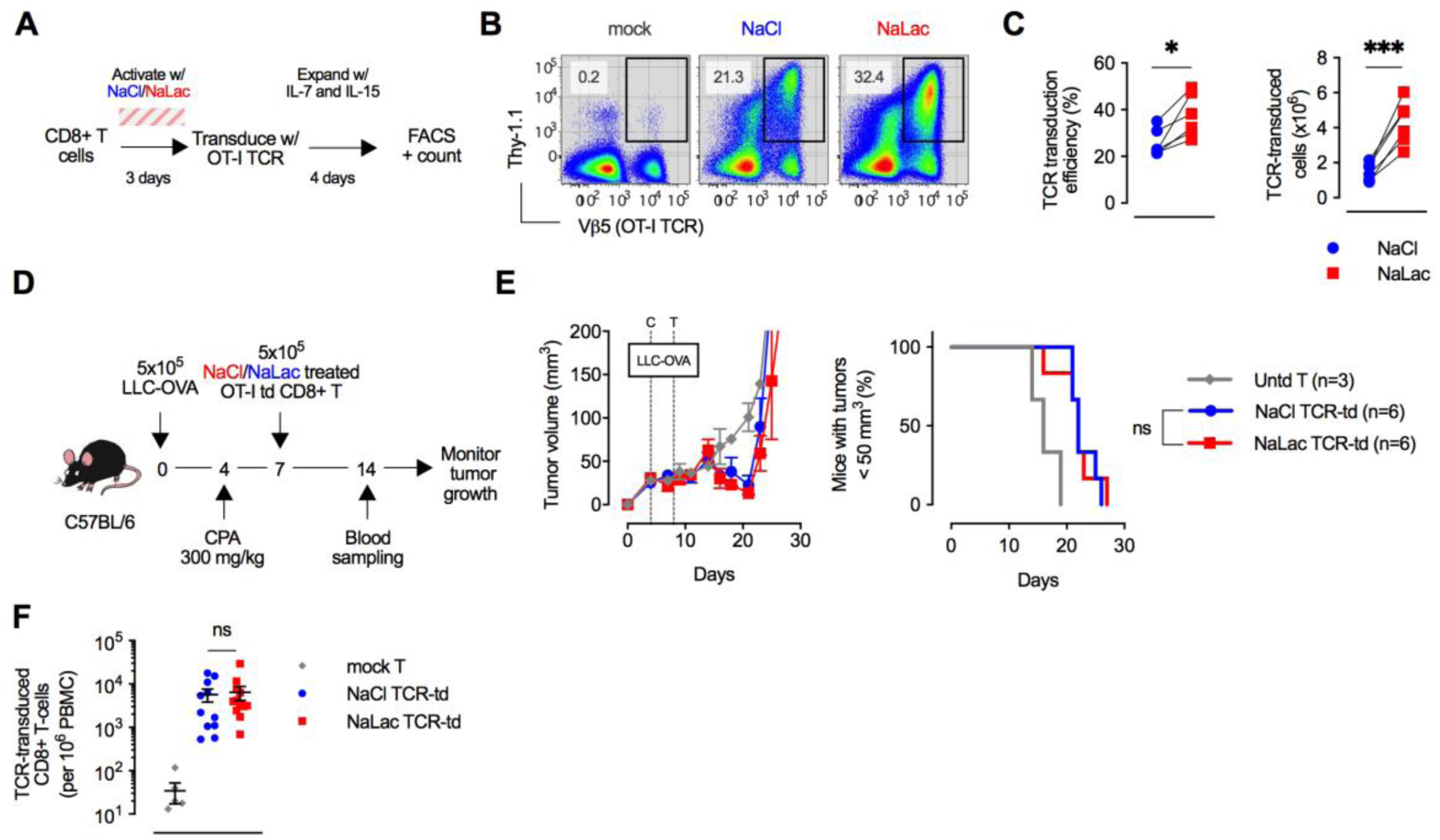
Sodium lactate enhances the *ex vivo* production of therapeutic CD8+ T cells. **A.** CD8+ T cells were purified from mouse splenocytes and activated with anti-CD3/CD28 beads and IL-2 for 3 days in the presence of either 40 mM NaCl or 40 mM NaLac. After activation, beads were removed, cells washed and counted before retroviral transduction with a vector encoding the OT-I TCR alpha (Vα2) and beta (Vβ5) chains and Thy-1.1 as a surface transduction marker. After transduction, cells were allowed to expand in the presence of IL-7 and IL-15 without further addition of NaCl or NaLac. Cell number and transduction efficiency were determined 4 days later. **B.** Representative flow cytometry plots of CD8+ T cells 4 days post-transduction with OT-I TCR. Transduced cells are Thy-1.1-positive (transduction marker) and Vβ5 positive (OT-I TCR beta chain). Comparison of mock-transduced cells and TCR-transduced cells previously activated in the presence of 40 mM NaCl or 40 mM NaLac. **C.** Frequency and absolute number of OT-I TCR-positive cells 4 days after transduction. Each pair represents an independent mouse donor (n=6). *** P<0.001, * P<0.05 two-tailed paired t-test. **D.** Experimental layout. C57BL/6 animals were inoculated with 5×10^5^ OVA-expressing Lewis Lung Carcinoma (LLC) cells and conditioned with 300 mg/kg CPA 4 days later. On day 7 tumour-bearing animals were injected with 5×10^5^ Thy-1.1+ Vβ5+ CD8+ T cells transduced with OT-I TCR as detailed in E, F, and G previously activated in the presence of 40 mM NaCl (NaCl TCR-td) or 40 mM NaLac (NaLac TCR-td) or mock-transduced CD8+ T cells. **E.** Plots showing mean and SEM tumour volume over time (left) and survival (right). ns, not significant, Log-rank (Mantel-Cox) survival test. **F.** Frequency of OT-I TCR-transduced CD8+ T cells in peripheral blood of tumour-bearing animals (as detailed in H and I) 7 days after T-cell adoptive transfer. Lines represent mean and SEM of n=12 mice (or n=6 for mock-transduced). Two-tailed unpaired t test.

Thus, exposure of CD8+ T cells to lactate during the initial stages of activation potentiates long-term cell expansion *ex vivo*. When combined with retroviral gene transfer, lactate-treatment enhances the production of therapeutic T cells by 3-fold without loss of potency upon transfer into tumour-bearing animals.

### Lactate alters CD8+ T cell metabolism

Lactate is transported across cell membranes via the proton-linked monocarboxylate transporters (MCT1, 2, 3, and 4)^30^ of which only MCT1 and MCT4 are expressed in CD8+ T cells^28,31^. While neither transporter is expressed in naïve CD8+ T cells, *ex vivo* activation causes MCT1 protein and mRNA to peak at 12 to 24 hours post activation while MCT4 levels peak later, between 48 and 72 hours (Supplementary Figure 4A and B). Activation of CD8+ T cells in the presence of an MCT1-specific inhibitor ablated cell proliferation at the low nanomolar range (Supplementary Figure 4C) suggesting a crucial role for monocarboxylate transport in early T-cell expansion.

Because metabolism and T-cell differentiation are linked^16,32^ we sought to characterise the CD8+ T-cell metabolome after activation in the presence of 40 mM sodium lactate (Figure 4A, Supplementary Figure 4D-G). Lactate treatment increased the frequency of glycolytic metabolites, namely glucose-6-phosphate (G6P), as well as 3-and 2-phosphoglyceric acid (3PGA, 2PGA). Pyruvate, the linking metabolite between glycolysis and the TCA cycle, was also increased. Within the TCA cycle, lactate increased the amount of succinate, and caused a reduction of the downstream metabolites fumarate and malate. Decreases in metabolites involved in amino acid metabolism and increases in metabolites linked to the amino sugar and pentose phosphate pathways were also observed (Supplementary Figure 4F). Interestingly, metabolic intermediates that are directly diverted into other routes do not accumulate (e.g., Glyceraldehyde 3-Phosphate).

**Figure 4.**
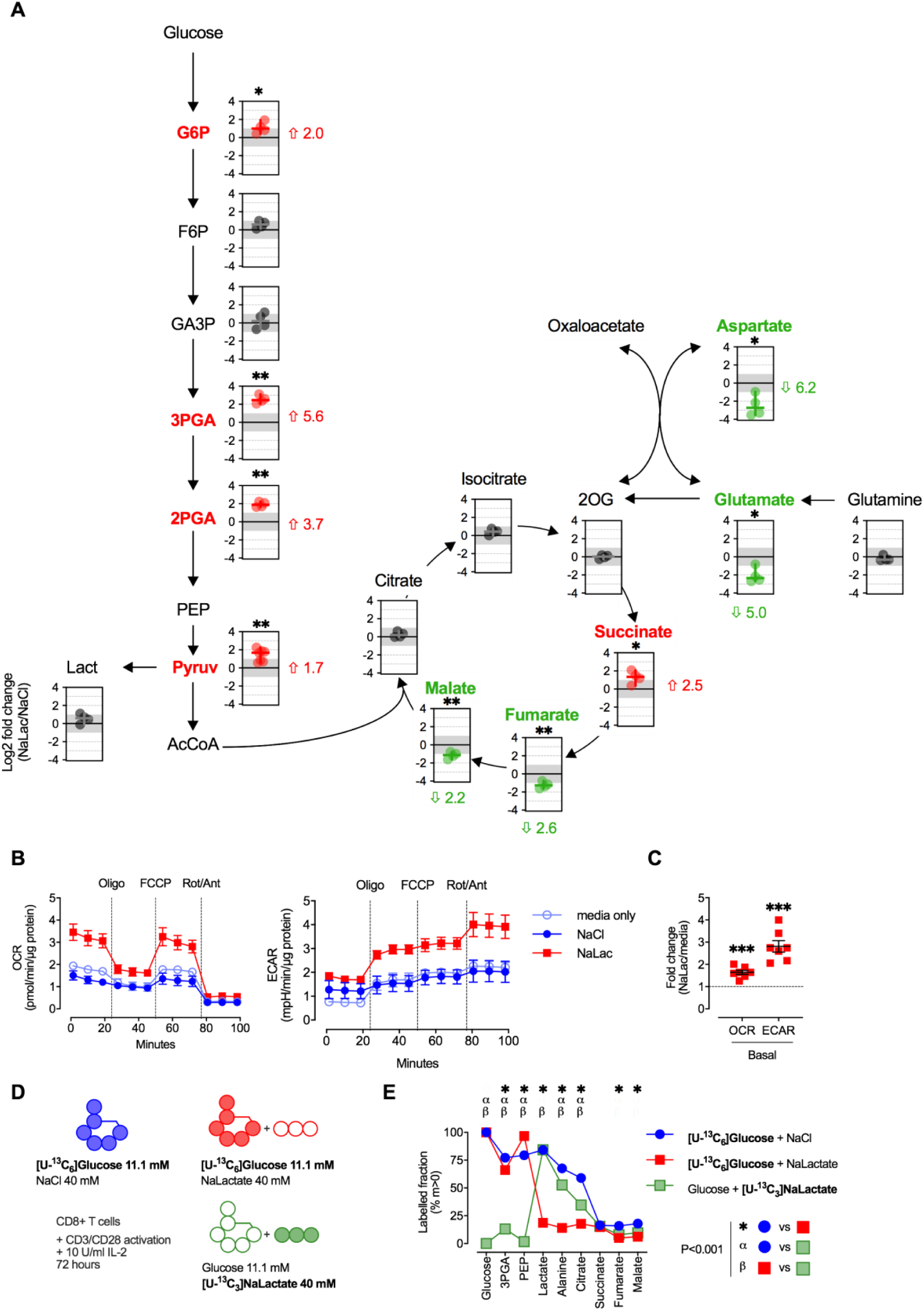
Lactate is utilized as a fuel by CD8+ T cells and alters the metabolic landscape. **A.** Metabolic changes after activation in the presence of lactate. Metabolites were extracted from mouse CD8+ T cells activated for 72 hours in the presence of 40 mM NaCl or NaLac and quantified by sugar phosphate and GC-MS analysis. Pyruvate levels were determined by colorimetric assay. Data shows log2 fold changes between NaLac and NaCl-treated samples of n=4 independent mouse donors. Red and green data points represent significantly increased or decreased metabolite frequency in NaLac-treated cells, respectively. Median fold-change next to each plot. ** P<0.01, *P<0.05, two-tailed ratio paired t-test. **B.** Oxygen consumption rate (OCR) and extracellular acidification rate (ECAR) of CD8+ T cells after 72 hours activation in the presence of 40 mM NaCl, 40 mM NaLac or media alone. After basal OCR and ECAR was measured cells were treated with 1 µM oligomycin (oligo), followed by 1.5 µM FCCP, and 100 nM rotenone and 1 µM antimycin A (Rot/Ant). OCR and ECAR are normalized to total protein content at end of assay. Data is the mean and SEM of n=4 independent mouse donors, each assayed as 4 technical replicates. **C.** Fold change in basal OCR and ECAR of CD8+ T cells activated for 72 hours in the presence of NaLac compared to media only of n=7 mouse donors. ***P<0.001, two-tailed ratio paired t-test. **D.** Substrate labelling experimental plan. Mouse CD8+ T cells were activated for 72 hours in the presence of 11.1 mM [U-^13^C_6_]Glucose and 40 mM NaCl, or 11.1 mM [U-^13^C_6_]Glucose and 40 mM NaLac, or 11.1 mM Glucose and 40 mM [U-^13^C_3_]NaLac. **E.** Proportion of metabolites labelled with at least one ^13^C (m>0) after incubation with 11.1 mM [U-^13^C_6_]Glucose or 40 mM [U-^13^C_3_]NaLac. Data is mean and SEM of n=4 independent mouse donors. *, *α, β* P<0.001, two-way ANOVA with Tukey’s multiple comparison test

Real time metabolic analyses revealed that CD8+ T cells activated in the presence of lactate have higher oxygen consumption and extracellular acidification rates, which are surrogate measurements of oxidative respiration and glycolysis, respectively (Figure 4B). Mitochondrial stress tests showed that lactate-treated T cells have higher basal and maximal respiration (Figure 4B). Basal respiration and glycolysis were increased by 1.5 and 2.8-fold, respectively (Figure 4C), demonstrating that high lactate levels can increase overall cellular metabolism of primary cytotoxic T cells in *ex vivo* culture.

To determine if lactate is taken up by CD8+ T cells and incorporated into carbon metabolism, and whether it displaces glucose as a carbon source, we activated T cells in the presence of [U-^13^C_6_]glucose (alone or in combination with lactate) or [U-^13^C_3_]lactate (Figure 4D). A progressive reduction in ^13^C enrichment in metabolites downstream of glucose was observed after 3 days of activation with [U-^13^C_6_]glucose alone, with higher labelling observed in glycolytic metabolites and lower in TCA cycle intermediates (Figure 4E). Addition of unlabelled lactate almost completely displaced the contribution of glucose to metabolites downstream of glycolysis, namely lactate, alanine, and citrate (Figure 4E). Activation in the presence of [U-^13^C_3_]lactate revealed little contribution of lactate to glycolytic metabolites and no conversion into glucose (Figure 4E). It did, however, show a significant incorporation of lactate into alanine, as well as the TCA metabolite citrate (Figure 4E). Lactate-derived ^13^C label was also incorporated at high frequency into other amino acids such as valine, and leucine (Supplementary Figure 4H).

This finding shows the unexpected importance of lactate as a carbon source for activated cytotoxic T cells and indicates that increased lactate levels produced in well-perfused tissues like skeletal muscle could have profound effects on the differentiation and efficacy of a cytotoxic T cell response.

### Sodium Lactate delays tumour growth in a CD8+ T cell-dependent manner *in vivo*

To determine how the lactate-induced increases in CD8+ T cell differentiation markers, cell numbers and cytotoxic efficacy might affect tumour growth *in vivo*, we performed infusions of sodium lactate into tumour-bearing animals at doses that result in plasma lactate levels similar to those seen during intensive exercise (approximately 10-20 mM). As can be seen in Supplementary Figure 5A, an intraperitoneal injection of a 2 g/kg dose of sodium lactate results in a 18 mM spike in serum lactate concentrations at 20 minutes post-injection. Following this dose, levels subside to 4 mM within 60 minutes; the expected time to reach baseline values from this magnitude of spike is approximately 180 minutes post-injection^33^. This dose was thus chosen as an approximation of the levels and persistence of rises in plasma lactate that occur following intense short-term periods of exercise.

In animals given daily doses of 2 g/kg sodium lactate, the effect was a marked and significant decrease in overall tumour growth after inoculations with I3TC on FVB animals (Figure 5A) as well as with the colon adenocarcinoma MC38 cell line on C57BL/6 animals (Figure 5C), with accompanying increases in tumour-bearing animal survival. Lower doses of lactate (0.5-1 g/kg) did not significantly alter tumour growth (Figure 5A), but a higher dose, 3 g/kg, also caused a significant reduction in tumour growth, with approximately the same efficacy as the 2 g/kg dose (Supplemental Figure 5B). As shown in Figure 5B, there were also significant changes in tumour infiltrating immune populations in animals treated with high doses of lactate, namely, increases in the frequency of total (CD3+) T cells, including both CD4+ and CD8+ T cells. The frequency of tumour-infiltrating NK cells was reduced in treated animals.

**Figure 5.**
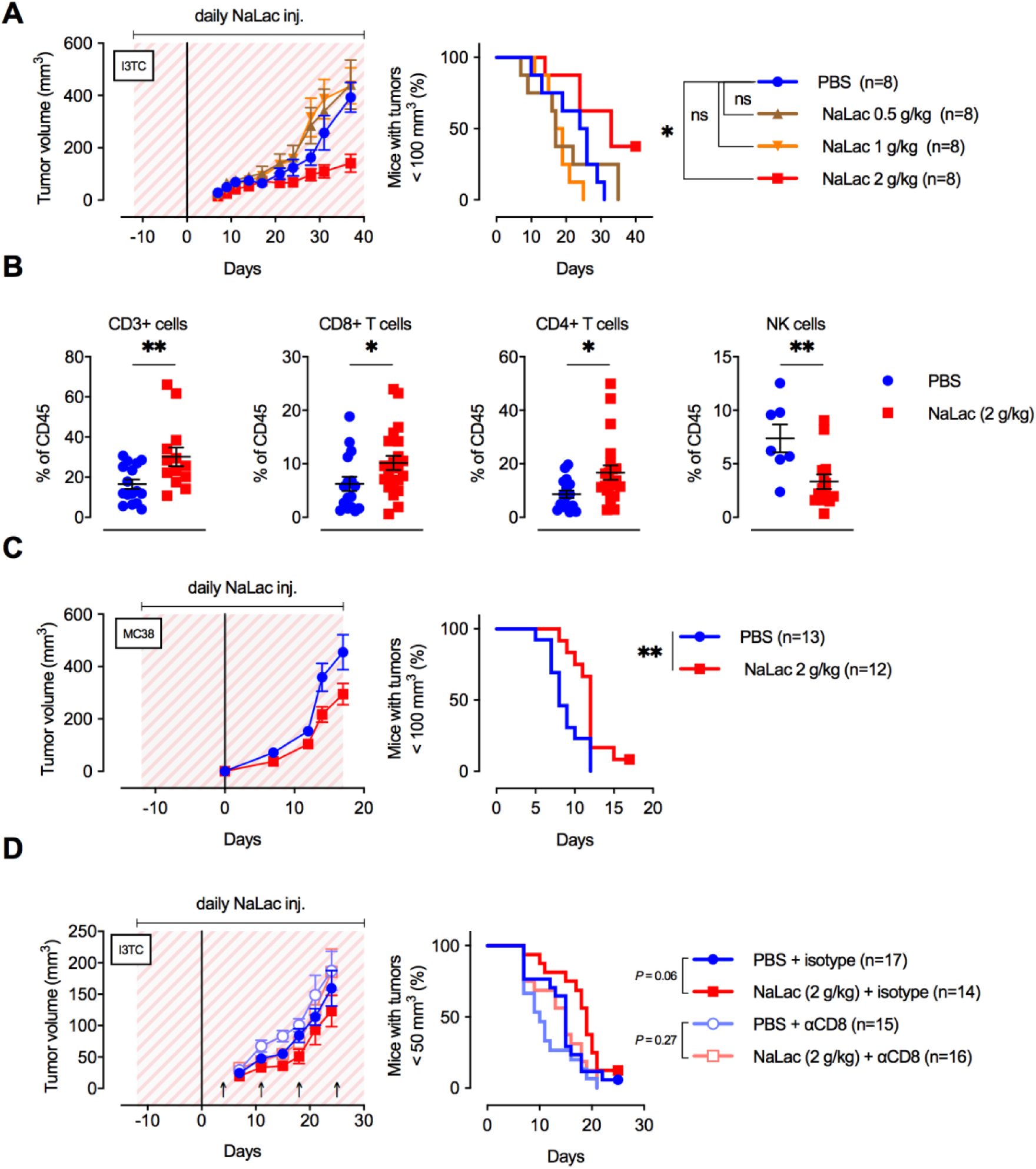
Daily administration of Sodium Lactate delays tumour growth. **A.** Daily doses of PBS or 0.5, 1 or 2 g/kg Sodium Lactate (NaLac) were administered i.p to FVB mice for 12 days before subcutaneous inoculation with 5×10^5^ cells of the MMTV-PyMT-derived breast cancer cell line I3TC. Daily sodium Lactate injections were continued throughout the experiment. Graphs show tumour volume (mean and SEM) over time and survival. * P <0.05, Log-rank (Mantel-Cox) survival test. **B.** Flow cytometric characterization of I3TC tumour infiltrating immune cell populations. ** P <0.01, * P <0.05, two-tailed t test. **C.** Daily doses of PBS or 2 g/kg NaLac was administered i.p to C57BL/6J mice for 12 days before subcutaneous inoculation with 5×10^5^ cells of the colon cancer cell line MC38. Injections were continued throughout the experiment. Graphs show tumour volume (mean and SEM) over time and survival. ** P <0.01, Log-rank (Mantel-Cox) survival test **D.** Same experimental setting as in (A) with or without weekly antibody-mediated depletion of CD8+ T cells (arrows). Graphs show tumour volume (mean and SEM) over time and survival. ns = not significant, Log-rank (Mantel-Cox) survival test.

To determine whether the effect of lactate infusion is dependent on CD8+ T cell-mediated effects on tumour growth, we depleted the cytotoxic T cell population via injection of an anti-CD8 antibody (Figure 5D) and observed that the net effect of lactate injection was reduced to statistical insignificance by the depletion of the CD8+ T cell population. This indicates that, at least in part, the effect of daily lactate injections on moderating tumour growth may be mediated by CD8+ T cells.

## Discussion

Here we show that exercise requires cytotoxic T cells to moderate tumour growth and progression. Recent studies suggest that exercise reduces cancer recurrence and mortality, and the effect of exercise on tumorigenic progression has now been documented in a range of animal models^34^. Previously proposed underlying mechanisms for the anti-neoplastic effects of exercise include effects on weight control and endocrine levels, as well as altered tumour vascularisation^35^ and immune function^36^. Exercise has been shown to reduce systemic inflammation^36^ and cause transient increases in circulating levels of immune cells^37^. Both CD8+ T cell and NK cell levels are increased in response to acute exercise. There is evidence that these immune cells populations exhibit an effector phenotype^38^ and an increased peripheral tissue homing capacity^39^, both important for anti-tumour activity.

Exercise is a multimodal stimulus. Systemic signalling from skeletal muscle activity to immune cell function has been attributed previously to skeletal muscle-derived myokines (e.g., IL-6 and OSM) and catecholamine release^36^. In a recent study, Hojman, et al., showed that an increase in systemic levels of epinephrine during exercise together with skeletal muscle IL-6 could increase NK cell recruitment during tumorigenesis^4^. However, in the mouse models employed here we found limited infiltration and no effect of exercise on the CD3- NK1.1+ population, relative to our observation of an increased frequency of CD3+ CD8+ T cells.

In evaluating the role metabolic flux plays in the process of modifying CD8+ T-cell function, we found that one key metabolite, lactate, has a highly unexpected function in the activation and differentiation of CD8+ T cells. A significant amount of recent data has shown that a wide range of mammalian tissues utilize lactate extensively during normal metabolism, and indeed, in many cases use it as a preferred carbon source^40-42^. Lactate generation is essential for the elimination of excessive pyruvate and for regeneration of NAD+ in order to sustain glycolytic flow, particularly when oxygen levels are limiting. Fast growing cell types, including malignant cells and activated lymphocytes, and metabolically active tissues such as exercising muscle, break down large amounts of glucose to meet energetic demands. As trafficking immune cells circulate they will often be exposed to high levels of lactate, in both loci of disease and in metabolically active tissues. Each lactate molecule is exported coupled with a proton, which is known to lead to a localized reduction in pH. However, healthy tissue microenvironments are very efficiently buffered and capable of maintaining a physiologically neutral pH even when large amounts of lactate are produced. In contrast, within poorly vascularised and fast-growing tumour masses, or in laboratory *ex vivo* cultures, lactate export results in a significant acidosis. Indeed, within the local micro-environment, tumour-derived lactate has been shown to inhibit the anti-tumour functions of T and NK cells due to intracellular acidification, in turn leading to NFAT suppression^27^. Likewise, lactate-associated acidity has been shown to impair CD8+ T cell motility^43^.

Thus, to mimic the activation of T cells in a lymphoid organ exposed to the physiologically buffered lactate generated by exercising muscle, we activated CD8+ T cells in the presence of a pH-neutral form of lactate: sodium lactate. Unlike its acid form, exposure to sodium lactate during the first 3 days of activation enhanced cytotoxic differentiation. Expression of both GzmB and IFNγ, two central proteins in anti-tumour function, was significantly enhanced after lactate exposure, and this was paired with a greater reduction in surface CD62L, a lymph node homing marker, and greater expression of the co-activator 4-1BB. Lactate had opposing effects on the levels of two co-inhibitors, CTLA-4 and PD-1, enhancing the former and reducing the latter. Despite this, lactate had an overall positive effect on cytotoxic activity, as measured by the ability of CD8+ T cells to kill tumour cells.

The supplied lactate was imported and used as a carbon source by CD8+ T cells and to a significant extent replaced glucose as the preferred substrate to generate amino acids and TCA cycle intermediates. The use of distally produced lactate as fuel for oxidative respiration has recently been reported in both malignant and non-malignant cell types^41,42^. This suggests that lactate can be a primary fuel used by CD8+ T cells, at least during early differentiation. Antigen presentation in lymph nodes is followed by extensive proliferation of antigen-specific T cells, all of which are engaging in glycolytic metabolism. Locally produced lactate originating from the primed T cells might, for example, function as a quorum sensing mechanism to cross-potentiate the early T cell response. In this context, exercise-derived lactate draining into an adjacent lymph node could act to boost a nascent T cell response.

We saw that the augmented effector differentiation phenotype caused by lactate was sustained in long term culture, but only if lactate exposure was limited to the first 3 days after activation. Short term, early lactate exposure also conditioned cells to proliferate more extensively. Our data indicate that at the levels we have employed, lactate has the capacity to significantly increase yields of differentiated cytotoxic T cells with increased cytotoxic efficacy and indicates that use of lactate supplementation could be an important new means to increase CD8+ T cell populations in adoptive T cell immunotherapy.

We have shown here the potential for treatment of tumour-bearing animals with lactate. The higher doses employed here have a significant effect on tumour growth. Further exploration of this effect will be needed to determine whether this would occur in other animal models of cancer and in human malignancy, but the non-toxic nature of sodium lactate makes it an intriguing candidate for further evaluation in cancer therapy.

## Materials and Methods

### Animals

All experiments and protocols were approved by the regional animal ethics committee of Northern Stockholm (dnr N78/15, N101/16). For the exercise experiments, female wild type (WT) FVB-mice, 6-7 weeks of age were purchased from Janvier Labs (France), allowed 7 days of acclimatization and housed 2 mice per cage in a temperature controlled room (20±2 °C) with dark-light cycles of 12 hours and access to food and water ad libitum. The running mice were allowed access to a running wheel (Med Associates Inc. ENV-044) and running patterns and distances were monitored wirelessly using appropriate software (Med Associates Inc. SOF-860 Wheel Manager and Med Associates Inc. SOF-861 Wheel Analysis). A control group with identical but locked wheels was included in the study to control for environmental enrichment.

For T cell purification, wild type donor and recipient C57BL/6J animals were purchased from Janvier Labs. TCR-transgenic OT-I mice (catalogue <u>003831</u>, The Jackson Laboratory) were crossed with mice bearing the CD45.1 congenic marker (catalogue <u>002014</u>, The Jackson Laboratory).

### Cell lines

HEK293 was a gift from Prof. N. Dantuma (Karolinska Institutet, Stockholm). MC38 was a gift from Dr. A. Palazón. EL4 was a gift from Prof. H. Stauss (UCL, London). B16-F10 and LLC were purchased from ATCC (<u>CRL-6475</u> and <u>CRL-1642</u>, respectively). I3TC was originally derived from the FVB MMTV-PyMT breast cancer model.

### Vectors

DNA encoding a codon-optimized polycistronic peptide composed of mouse Thy-1.1 (<u>AAR17087.1</u>) and the alpha and beta chains of the mouse OT-I TCR interspersed with furin (RAKR) and picornavirus P2A (GSGATNFSLLKQAGDVEENPGP) cleavage sequences was synthesized by Gene Art (Thermo Fisher). Cell surface localization peptides were incorporated into the Thy-1.1 and TCR sequences. The ORF was cloned into the gamma retroviral vector pMP71, a gift from Christopher Baum (MHH, Hannover). DNA encoding a codon-optimized polycistronic peptide composed of chicken ovalbumin (OVA; P01012.2), eGFP (ABG78037.1) and neomycin phosphotransferase (NeoR; BAD00047.1) interspersed with P2A and furin cleavage sites was synthesized by Gene Art (Thermo Fisher) and cloned under control of the SV40 promoter in the transposon vector pT2/BH, a gift from Perry Hackett (Addgene plasmid #26556). pCMV-SB11 encoding the sleeping beauty transposase was a gift from Perry Hackett (Addgene plasmid #26552).

### Generation of ovalbumin-expressing cell lines

B16-F10, LLC and EL4 cells were co-transfected with the transposon vector pT2 encoding OVA, eGFP and neomycin phosphotransferase and the vector encoding transposase SB11. Three days later 400 mg/ml G418 (Gibco, <u>10131035</u>) was added to culture media to select for transgene-expressing cells. Successful integration was confirmed by analyzing eGFP fluorescence by flow cytometry. Limiting dilution was used to derive monoclonal OVA-expressing lines for each cell line. OVA presentation was confirmed by flow cytometry using a PE-labeled antibody against surface SIINFEKL bound to H-2Kb (clone 25-D1.16, BioLegend).

### Tumour single cell suspension

Tumour samples were dissected and minced with scissors, resuspended in 2 ml Digestion buffer composed of HBSS with 2 mg/ml Collagenase A (<u>50-100-3278</u>, Fischer Scientific) and 10 µg/ml DNase (<u>D5025</u>, Sigma-Aldrich), dissociated with a gentleMACS Dissociator (Miltenyi Biotec, <u>130-093-235</u>) and incubated 30 min at 37°C. 10 ml PBS with 10 % FCS (buffer A) was added and the suspension was passed through a 100 µm cell strainer (BD Biosciences). The resulting single cell suspension was pelleted and resuspended in 1 ml ACK buffer (0.15 M NH_4_Cl, 10 mM KHC0_3_, 0.1 mM EDTA in distilled H_2_O) and incubated on ice for 5 minutes. 10 ml of buffer A was added to stop the reaction. The suspension was passed through a 40 µm cell strainer (BD Biosciences), pelleted and resuspended in 500 µL of cold PBS. Single cell suspensions were subsequently subject to cell phenotyping by flow cytometry as described below.

### Mouse CD8+ T-cell purification, activation and expansion

Spleens were harvested from 8-12 week old C57BL/6J mice, mashed over a 40 µm cell strainer (VWR, <u>10199-654</u>), and CD8+ T-cells purified by positive magnetic bead selection (Miltenyi Biotec, <u>130-117-044</u>) according to manufacturer’s instructions. Purified CD8+ T-cells were counted and cell diameter measured using a Moxi Z mini automated counter (Orflo, <u>MXZ001</u>). 1×10^6^ (24-well plate) or 5×10^5^ (48-well plate) CD8+ T-cells were activated with Dynabeads Mouse T-Activator CD3/CD28 (Thermo Fisher, <u>11456D</u>) in a 1:1 beat to T-cell ratio and cultured in 2 ml (24WP) or 1 ml (48WP) RPMI 1640 (Thermo Fisher, <u>21875</u>) supplemented with 10% Fetal Bovine Serum, (Thermo Fisher, <u>10270-106</u>), 50 µM 2-mercaptoethanol (Thermo Fisher, <u>21985023</u>), 10 U/ml penicillin-streptomycin (Thermo Fisher, <u>15140122</u>), and 10 U/ml recombinant human IL-2 (Sigma, <u>11011456001</u>), and incubated at 37°C for 3 days in a humidified CO_2_ incubator.

For long-term expansion after 3 days, dynabeads were removed using a DynaMag-2 Magnet (Thermo Fisher, <u>12321D</u>), washed with PBS, counted, and 1×10^6^ T-cells resuspended in 2 ml (24WP) or 5×10^5^ T-cells resuspended in 1 ml (48WP) in fresh media supplemented with 10 U/ml IL-2 or 10 µg/ml recombinant human IL-7 (RnD Systems, <u>207-IL-005</u>) together with 10 µg/ml recombinant human IL-15 (RnD Systems, <u>247-IL-005</u>) and incubated as described above. Media supplemented with cytokines was added every 2-3 days in order to maintain cell densities below 2×10^6^/ml. Cells were re-stimulated at day 7 and day 14 with CD3/CD28 Dynabeads as described above.

### Human CD8+ T-cell purification, activation and expansion

Peripheral blood mononuclear cells (PBMCs) were harvested from standard buffy coat preparations of healthy donors, aged 20 to 40 years old, obtained from the Department of Transfusion Medicine at Karolinska University Hospital and processed no later than 4 hours after collection. PBMCs were isolated by gradient centrifugation using Histopaque-1077 (Sigma, <u>10771</u>) and enriched for Naïve CD8+ T-cells by magnetic bead selection, according to manufacturer’s instructions (Miltenyi Biotec, <u>130-093-244</u>).

Purified naïve CD8+T-lymphocytes were counted and 5×10^5^ (48-well plate) or 1×10^5^ (96-well plate) naïve CD8+ T-cells were activated with Dynabeads Human T-Activator CD3/CD28 (Thermo Fisher, <u>11132D</u>) in a 1:1 beat to T-cell ratio and cultured in 1 ml (48WP) or 200µl (96 WP) RPMI 1640 (Thermo Fisher, <u>21875</u>) supplemented with 10% Fetal Bovine Serum, (Thermo Fisher, <u>10270-106</u>), 10 U/ml penicillin-streptomycin (Thermo Fisher, <u>15140122</u>), and 30 U/ml recombinant human IL-2 (Sigma, <u>11011456001</u>), and incubated at 37°C for 4 days in a humidified CO_2_ incubator.

For long-term expansion after 4 days, Dynabeads were removed using a DynaMag-2 Magnet (Thermo Fisher, <u>12321D</u>), washed with PBS, counted, and resuspended in 1 ml (48WP) or 200µl (96 WP) of fresh media supplemented with 30 U/ml IL-2 and incubated as described above. Media supplemented with cytokines was added every 2-3 days in order to maintain cell densities below 2×10^6^/ml. Cells were counted again at day 7 and maintained in culture under the above described conditions until day 14.

### CD8+ T cell *ex vivo* activation

Sodium L-Lactate (Sigma, <u>L7022</u>), Sodium D-Lactate (Sigma, <u>71716</u>), Sodium L-Pyruvate (Sigma, <u>P2256</u>), L-Lactic Acid (Sigma, <u>L1750</u>) and Sodium Chloride (Sigma, <u>S5886</u>) were prepared as 10x concentrated solutions in complete media. Compounds were added to T-cells at the point of activation (day 0) minutes before addition of CD3/CD28 Dynabeads. Sodium Chloride or plain media was used as control. For long-term expansion, after bead removal at day 3, cells are cultured in complete media supplemented with cytokines without further addition of compounds.

### Cell proliferation analysis

After magnetic bead purification, CD8+ T-cells were loaded with 5 µM CellTrace CFSE or CellTrace Violet (CTV) dyes (Thermo Fisher, <u>C34554</u> and <u>C34557</u>, respectively) according to manufacturer’s instructions. CFSE or CTV dilution was determined by flow cytometry.

### Cell phenotyping by flow cytometry

Single-cell suspensions were washed and stained with Fixable Near-IR Dead Cell Stain Kit (Thermo Fisher, <u>L10119</u>) followed by staining of extracellular antigens with fluorochrome labelled antibodies. The Fixation/Permeabilization Solution Kit (BD Biosciences, <u>554714</u>) was used for exposing cytoplasmic antigens. The Transcription Factor Buffer Set (BD Biosciences, <u>562574</u>) was used for exposing nuclear antigens. Fluorochrome-labelled antibodies against mouse antigens CD44 (clone IM7), CD45.1 (clone A20), CD45.2 (clone 104), CD8 (clone 53-6.7), CTLA-4 (clone UC10-4F10-11), LAG3 (clone C9B7W), PD-1 (clone J43), and Vα2 TCR (clone B20.1) were purchased from BD Biosciences, CD27 (clone LG.3A10), CD28 (clone 37.51), CD62L (clone MEL-14), and ICOS (clone C398.4A) were purchased from BioLegend, and 4-1BB (clone 17B5), CD127 (clone A7R34), CD25 (clone PC61.5), Eomes (clone Dan11mag), T-bet (clone eBio4B10), Thy-1.1 (clone HIS51), Vβ5.1/5.2 TCR (clone MR9-4), and anti-human/mouse Granzyme B (clone GB12) was purchased from Thermo Fisher. Antibodies against human CD62L (clone DREG-56), CCR7 (clone 3D12), CD45RA (clone HI100), CD45RO (clone UCHL1), CD8 (clone SK1), CD44 (clone IM7) were purchased from BD Biosciences. Cell counting was performed with CountBright Absolute Counting Beads (Thermo Fisher, <u>C36950</u>). Samples were processed in a FACSCanto II flow cytometer (BD Biosciences). Data analysis was performed with FlowJo, version 8.8.7 (Tree Star).

### Western blotting and real time RT-PCR

Mouse CD8+ T cells were activated ex vivo and cells harvested at 0, 6, 12, 24, 48 and 72 hours post-activation. For each time point, total protein was extracted using RIPA buffer (R0278, Sigma) and total RNA extracted using RNeasy Mini Kit (74104, Qiagen). For each sample, 15 µg of total protein were separated in SDS-PAGE, blotted onto a PVDF membrane and probed with antibodies against MCT1 (sc-50325, Santa Cruz), MCT4 (sc-50329, Santa Cruz), GzmB (4275, Cell Signaling Technologies) and PPIB (A7713, Antibodies Online) and detected using infra-red labeled secondary antibodies in an Odyssey imaging system (LI-COR). REVERT Total protein stain (926-11010, Li-COR) was used for lane normalization. Two micrograms of total RNA were reverse transcribed using iScript cDNA synthesis kit (BioRad) in a total volume of 20 μl. Real-time RT-PCR was used for mRNA quantification (7500 Fast Real-Time PCR system, Applied Biosystems Inc., Foster City, California, USA). All primers were designed to cover exon-exon boundaries to avoid amplification of genomic DNA. SYBR-green predesigned primers used to detect mouse *Slc16a1* (forward: ACTTGCCAATCATAGTCAGAGC, reverse: CGCAGCTTCTTTCTGTAACAC), *Slc16a3* (forward: GACGCTTGTTGAAGTATCGATTG, reverse: GCATTATCCAGATCTACCTCACC), *GzmB* (forward: CTGCTAAAGCTGAAGAGTAAG, reverse: TAGCGTGTTTGAGTATTTGC), and *Hprt* (forward: TGACACTGGCAAAACAATGCA, reverse: GGTCCTTTTCACCAGCAAGCT) were purchased from Sigma. All reactions were performed in 96-well MicroAmp Optical plates in duplicates. The total reaction volume was 15 µl, containing 5 µl sample cDNA, 0.4 mM of each primer forward and SYBR Green PCR Master Mix (4309155E, Applied Biosystems Ins.) All quantification reactions were controlled with a melting curve and primer efficiency was tested with standard curves, and did not differ between the primer pairs. Gene expression data was normalized to *Hprt* levels.

### Immunohistochemistry characterisation of primary tumours

Mammary gland sections were deparaffinized with Tissue-Tek Tissue-Clear (Sakura, Japan) and rehydrated with graded ethanol. Antigen retrieval was performed either with Proteinase K (for F4/80) or boiling the sections in a high pH antigen retrieval buffer (Dako Target retrieval Solution, pH 9, Dako, Denmark) (for Podocalyxin, CD3, Granzyme B) or low pH antigen retrieval buffer (Dako Target retrieval Solution, pH 6, Dako, Denmark) (for PCNA). Endogenous peroxidase activity was blocked using 3 % H_2_O_2_. Nonspecific protein interactions were blocked using 20 % goat serum in PBS-T. Primary antibodies were used in lungs and mammary glands against CD3 (#ab5690, Abcam, UK) to assess lymphocytic infiltration, F4/80 (#MCA497, Abd Serotec, Germany) to assess macrophage density, podocalyxin (#AF1556, R&D Systems, Germany) to assess capillary density, PCNA (#M0879, Dako, Denmark) to identify proliferating cells, and Granzyme B (#ab4059, Abcam, UK) to identify effector lymphocytes. Subsequently the sections were incubated with a suitable biotin conjugated secondary IgG antibody. All sections were then incubated with an Avidin-Biotin peroxidase Complex (ABC) kit (#PK6100, Vector, USA) and the peroxidase was stained with 3,3’-Diaminobenzidine(DAB) kit (#SK4100, Vector, USA). Nuclei were stained with hematoxylin and sections were mounted with Faramount Aqueous Mounting Medium, (Dako, Denmark). Specific stainings were quantified using Image J software (National Institute of Health, USA)

### Histological characterisation

Mammary glands were stained with H&E with a standard protocol and histologically scored for tumour stage in accordance with ^19^, ranging from hyperplasia (score 1), adenoma/mammary intraepithelial neoplasia (adenoma/MIN) (score 2), early invasive carcinoma (score 3) to late invasive carcinoma (score 4). The highest staging value of each specimen was used.

### Metabolite extraction

For the metabolite analysis 5×10^6^ cells were harvested and washed 3 times with PBS before adding 800µL of ice-cold extraction mixture consisting of chloroform:MeOH:H_2_O (1:3:1), containing 500 pg/µL of 2-Deoxy-D-glucose 6-phosphate and 70 pg/µL of myristic acid-13C3 were added to each sample. The metabolites were extracted using a mixer mill set to a frequency 30 Hz for 2 min, with 1 tungsten carbide bead added to each tube. Obtained extracts were centrifuged at 14000 rpm for 10 min. The collected supernatants were divided for GCMS and Sugar-P analysis. 300 µL of the supernatant was transferred into GC and LC-vial respectively, the supernatants were evaporated until dryness using a SpeedVac.

### Sugar phosphate analysis

For derivatization, dried samples were dissolved in 20µl of methoxylamine and incubated on a heat block at 60°C, 30 min. After overnight incubation at room temperature, 12µl of 1-Methylimidazol and 6 µl of propionic acid anhydride were added and heated at 37°C for 30 minutes. The reaction mixture was then evaporated to dryness by N2 gas. Prior to LC-MS analysis, derivatized metabolites were dissolved in 100µl of aqueous 0.1% formic acid.

Quantitative analysis was done by combined ultra-high-performance liquid chromatography-electrospray ionization-triple quadrupole-tandem mass spectrometry (UHPLC-ESI-QqQ-MS/MS) in dynamic multiple-reaction-monitoring (MRM) mode. An Agilent 6495 UHPLC chromatograph equipped with a Waters Acquity BEH 1.7 µm, 2.1 x 100mm column (Waters Corporation, Milford, USA) coupled to a QqQ-MS/MS (Agilent Technologies, Atlanta, GA, USA) was used. The washing solution, for the autosampler syringe and injection needle, was 90% MeOH with 1% HCOOH. The mobile phase consisted of A, 2% HCOOH and B, MeOH with 2% HCOOH. The gradient was 0% B for 1 min followed by linear gradients from 0.1 to 30% from 1 to 3 min then 30 to 40% B from 3 to 6 min, hold at 40% B from 6 to 10 min, followed by 40 to 70% B from 10 to 12.5 min, hold at 70% B from 12.5 to 15 min, and thereafter 70 to 99% B from 15 to 17.5 min. B was held at 99% for 0.5 min, and thereafter the column was re-equilibrated to 0% B. The flow rate was 0.65 mL min-1 during equilibration and 0.5 mL min-1 during the chromatographic runs. The column was heated to 40 °C, and injection volumes were 1 μL. The mass spectrometer was operated in negative ESI mode with gas temperature 230°C; gas flow 12 L min-1; nebulizer pressure 20 psi; sheath gas temperature 400°C; sheath gas flow 12 L min-1; capillary voltage 4000 V (neg); nozzle voltage 500 V; iFunnel high pressure RF 150 V; iFunnel low pressure RF 60 V. The fragmentor voltage 380 V and cell acceleration voltage 5 V. For a list of MRM transitions see Suppl. Data Table S1. Data were processed using MassHunter Qualitative Analysis and Quantitative Analysis (QqQ; Agilent Technologies, Atlanta, GA, USA) and Excel (Microsoft, Redmond, Washington, USA) software.

### GCMS analysis

The GCMS samples were spiked of with 1050 pg of each GCMS internal standard before evaporation. Derivatization was performed according to Gullberg et al ^44^. In detail, 10µL of methoxyamine (15 µg/µL in pyridine) was added to the dry sample that was shaken vigorously for 10 minutes before left to react in room temperature. After 16 hours 10 µL of MSTFA was added, the sample was shaken and left to react for 1 hour in room temperature. 10 µL of methyl stearate (1050 pg/µL in heptane) was added before analysis. One µL of the derivatized sample was injected by an Agilent 7693 autosampler, in splitless mode into an Agilent 7890A gas chromatograph equipped with a multimode inlet (MMI) and 10 m x 0.18 mm fused silica capillary column with a chemically bonded 0.18 μm DB 5-MS UI stationary phase (J&W Scientific). The injector temperature was 260 °C. The carrier gas flow rate through the column was 1 ml min-1, the column temperature was held at 70 °C for 2 minutes, then increased by 40 °C min-1 to 320 °C and held there for 2 min. The column effluent is introduced into the electron impact (EI) ion source of an Agilent 7000C QQQ mass spectrometer. The thermal AUX 2 (transfer line) and the ion source temperatures were 250 °C and 230 °C, respectively. Ions were generated by a 70 eV electron beam at an emission current of 35 µA and analyzed in dMRM-mode. The solvent delay was set to 2 minutes. For a list of MRM transitions see Suppl. Data Table S2. Data were processed using MassHunter Qualitative Analysis and Quantitative Analysis (QqQ; Agilent Technologies, Atlanta, GA, USA) and Excel (Microsoft, Redmond, Washington, USA) software.

### Isotopic labelling, metabolite extraction and GC/MS analysis

Mouse CD8+ T cells were activated with CD3/CD28 beads and 10 U/ml IL-2 in glucose-free RPMI supplemented with 2mM glutamine, 10% FBS, penicillin/streptomycin and 2-mercaptoethanol, and either i) 11 mM [U-^13^C_6_] glucose, ii) 11mM [U-^13^C_6_] glucose and 40 mM lactate, or iii) 11 mM glucose and 40 mM [U-^13^C_3_] sodium lactate, and isotopic labelling was performed for 72 hours. Sodium chloride (40mM) was added to i) to maintain osmotic concentration. At the conclusion of the labelling experiment, cells were washed with PBS, and metabolic activity quenched by freezing samples in dry ice and ethanol, and stored at −80°C. Metabolites were extracted by addition of 600 μl ice-cold 1:1 (vol/vol) methanol/water (containing 1 nmol scyllo-Inositol as internal standard) to the cell pellets, samples were transferred to a chilled microcentrifuge tube containing 300μl chloroform and 600μl methanol (1500 μl total, in 3:1:1 vol/vol methanol/water/chloroform). Samples were sonicated in a water bath for 8 min at 4°C, and centrifuged (13000 rpm) for 10 min at 4°C. The supernatant containing the extract was transferred to a new tube for evaporation in a speed-vacuum centrifuge, resuspended in 3:3:1 (vol/vol/vol) methanol/water/chloroform (350μl total) to phase separate polar metabolites (upper aqueous phase) from apolar metabolites (lower organic phase), and centrifuged. The aqueous phase was transferred to a new tube for evaporation in a speed-vacuum centrifuge, washed with 60 μl methanol, dried again, and derivatized by methoximation (20μl of 20 mg/ml methoxyamine in pyridine, RT overnight) and trimethylsilylation (20μl of N,O-bis(trimetylsilyl)trifluoroacetamide + 1% trimethylchlorosilane) (Sigma, 33148) for ≥ 1 h. GC/MS analysis was performed using an Agilent 7890B-5977A system equipped with a 30 m + 10 m × 0.25 mm DB-5MS + DG column (Agilent J&W) connected to an MS operating in electron-impact ionization (EI) mode. One microliter was injected in splitless mode at 270 °C, with a helium carrier gas. The GC oven temperature was held at 70°C for 2 min and subsequently increased to 295 °C at 12.5 °C/min, then to 320 °C at 25 °C/min (held for 3 min). MassHunter Workstation (B.06.00 SP01, Agilent Technologies) was used for metabolite identification by comparison of retention times, mass spectra and responses of known amounts of authentic standards. Metabolite abundance and mass isotopologue distributions (MID) with correction of natural 13C abundance were determined by integrating the appropriate ion fragments using GAVIN.

### Quantification of interferon-gamma secretion

Soluble Interferon-gamma (IFN-gamma) was quantified in culture media 3 days after CD8+ T-cell activation using the Mouse IFN-gamma Quantikine ELISA Kit (RnD Systems, <u>MIF00</u>).

### Cytotoxicity assay

OVA-expressing GFP-positive EL4 (EL4-GFP-OVA) cells were mixed with their respective parent cell line (EL4) in a 1:1 ratio in round-bottom 96-well plates. OVA-specific OT-I CD8+ T-cells were mixed with EL4 cells at effector to target ratios ranging from 20:1 to 1:50. Cytotoxicity was assessed by flow cytometry 24 hours later. The ratio of GFP-positive events (target) to GFP-negative events in each test co-culture was divided by the ratio from cultures without addition of OT-I cells to calculate specific cytotoxicity.

### Oxygen consumption rate and extracellular acidification rate measurements

CD8+ T cells activated for 3 days were washed and oxygen consumption rate (OCR) and extracellular acidification rate (ECAR) in a Seahorse Extracellular Flux Analyzer XF96 (Agilent). 5×10^5^ CD8+ T-cells were plated onto poly-D-lysine coated wells and assayed in XF RPMI medium (Agilent) pH 7.4 supplemented with 10 mM glucose, 1 mM sodium pyruvate and 2 mM glutamine. During the assay wells were sequentially injected with 1 µM oligomycin (Sigma), 1.5 µM FCCP (Sigma) and 100 nM rotenone (Sigma) + 1 µM antimycin A (Sigma).

### Retroviral transduction

Sub-confluent HEK293 cultures were transfected with OT-I vectors and helper vector pCL-Eco, a gift from Inder Verma (Addgene plasmid #12371). Supernatant media containing retroviral particles was harvested 48 hours after transfection and used fresh or stored at −80°C. Retroviral supernatants were spun onto Retronectin-coated wells (Takara) and replaced with CD8+ T-cells activated for 3 days in fresh RPMI supplemented with 10 U/ml IL-2. Fresh media was added every 2-3 days. Transduction efficiency was assessed by measuring surface expression of the transduction marker Thy-1.1, V*α*2 TCR chain, and V*β*5 TCR chain by flow cytometry.

### *In vivo* tumour growth

Four week old PyMT+ females of the MMTV-PyMT strain were housed individually in a temperature controlled room (20±2 °C) with 12 hour dark-light cycles and food and water ad libitum. The mice were allowed free access to a running wheel (Med Associates Inc. ENV-044, St Albans, USA). Running patterns and distance were monitored wirelessly using appropriate software (Med Associates Inc. SOF-860 Wheel Manager and Med Associates Inc. SOF-861 Wheel Analysis). A control group with identical but locked wheels was included in the study to control for environmental enrichment. Animal body weight was monitored weekly throughout the experiment and mammary gland tumours measured twice weekly using calipers. Tumour volumes (V) were approximated using the formula V = (π/6) * l * w * h. Experiments were terminated when the animals reached 12 weeks of age. FVB WT animals were introduced to running or locked wheels and allowed to acclimate to running for two weeks prior to injections of 5×10^5^ tumour cells of the I3TC cell line. The cells were trypsinized, washed once with PBS and suspended in 100 µL sterile PBS and injected subcutaneous (s.c) into running and non-running FVB WT mice. Animal body weight was monitored throughout the experiment. Tumours size was measured as previously described. Experiments were terminated 8 weeks after tumour cell injections. Upon termination of the experiments, tumour specimens were surgically removed and fixed in 4% paraformaldehyde (Sigma, <u>F8775</u>) or dissociated for single cells suspension.

For CD8 depletion, 200 µg of antibodies (anti-mCD8α clone 53-6.72, BioXCell or Rat IgG2a Isotype control, clone 2A3, BioXCell) was injected i.p at day 4 post tumour cell injections and repeated weekly throughout the experiment.

To evaluate the effect of lactate on tumour growth in vivo, FVB and C57/Bl6J WT animals were subject to daily intraperitoneal (i.p) Sodium L-Lactate (Sigma, <u>L7022</u>) injections (0.5, 1, 2, or 3 g/kg) for 12 days prior to injections of 5×10^5^ tumour cells of the I3TC or MC38 cell line. Tumour growth was monitored as previously described and daily injections of Sodium L-Lactate was continued throughout the experiment.

For T-cell adoptive transfer, 8 to 12-weeks old female C57BL/6J were inoculated sub-cutaneously with 5×10^5^ B16-F10-OVA or LLC-OVA and conditioned 4 days later with peritoneal injection of 300 mg/kg cyclophosphamide (Sigma, <u>C0768</u>) in PBS. At day 8, 1×10^6^ or 1×10^7^ transgenic OT-I or 5×10^5^ OT-I-transduced CD8+ T-cells were injected peritoneally. Tumour volume was measured every 2-3 days with electronic calipers until day 60. At day 15, 7 days after OT-I cell transfer, peripheral blood was collected from tail vein, red blood cells lysed with water, and leukocyte cell suspension analyzed by flow cytometry. To ascertain transient levels of sodium lactate in plasma induced by lactate injection, 2g/kg of Sodium L-Lactate was administered i.p to FVB and C57Bl6 animals (n=1), systemic levels of lactate were monitored through tail vein bleedings pre-injection, and at 5,10, 20, 40, and 60 minutes post-injection (Accutrend®Plus).

### Statistics

Statistical analyses were performed with Prism 7 version 7.0 (GraphPad). Statistical tests and number of replicates are stated in figure legends.

### Study approval

All animal experiments were approved by the regional animal ethics Committee of Northern Stockholm, Sweden (dnr N78/15 and N101/16). The Stockholm Regional Ethical Review Board does not require an ethical permission for the use of non-identified healthy donor samples.

## Acknowledgements

The authors acknowledge Dr. David Macias (University of Cambridge) for advice on osmolality measurements, Carolin Lindholm (Karolinska Institutet) for expertise in immunohistochemistry, Prof. Kristian Pietras (Lunds Universitet) for sharing the MMTV-PyMT mouse strain on the FVB background, Dr. Carina Strell (Karolinska Institutet) for sharing the I3TC cell line, and Prof. Ananda Goldrath and Prof. Christer Höög for discussions. Metabolomics experiments were performed in the Metabolomics STP at The Francis Crick Institute, which receives its core funding from Cancer Research UK (FC001999), the UK Medical Research Council (FC001999), and the Wellcome Trust (FC001999). The work was funded by the Swedish Medical Research Council (Vetenskapsrådet), the Swedish Cancer Fund (Cancerfonden), the Swedish Children’s Cancer Fund (Barncancerfonden), and a Principal Research Fellowship to RSJ from the Wellcome Trust. We would like to thank James MacRae and James Ellis for help with the GC-MS analysis.

---

**Supplementary Figure 1.**
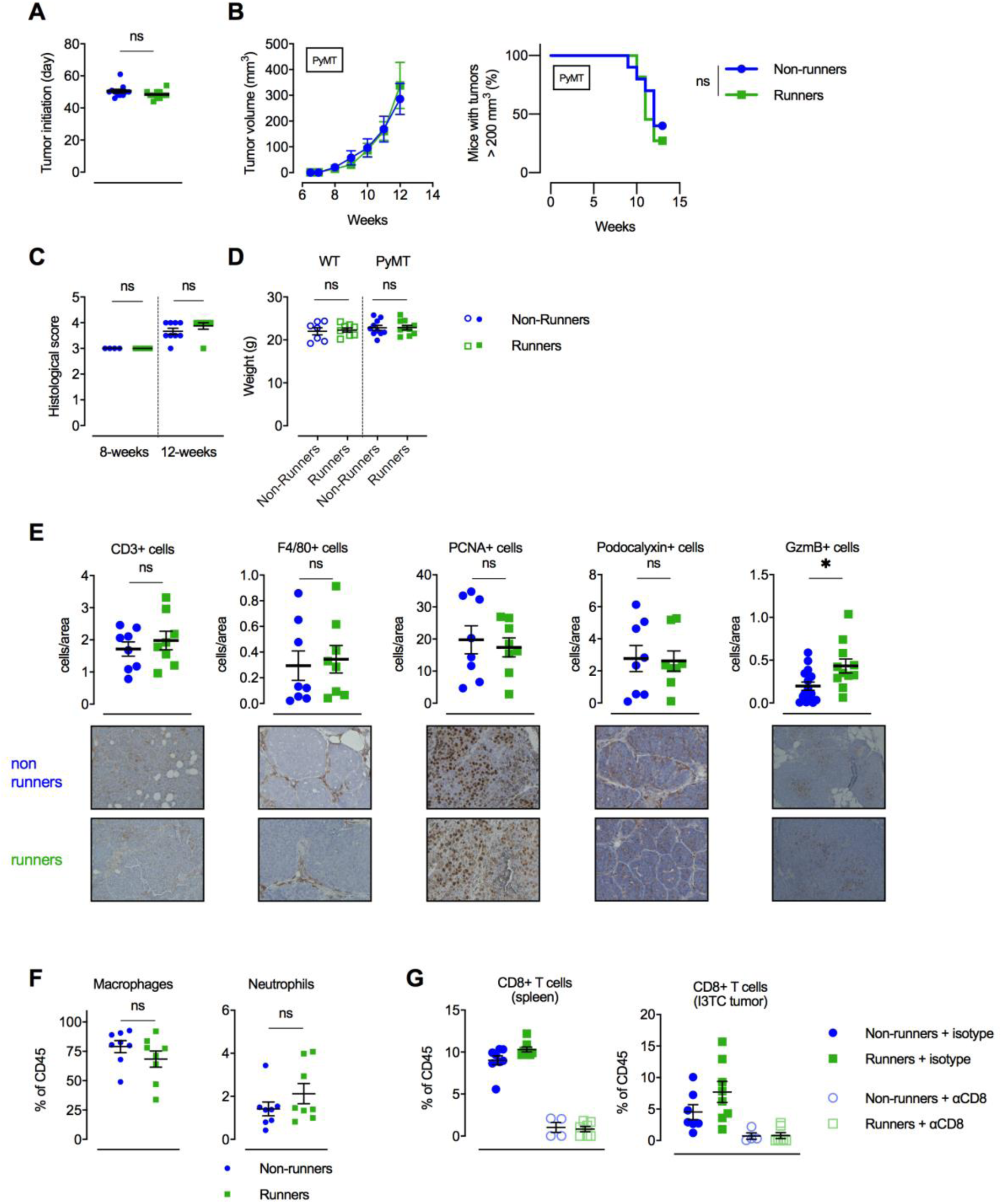
Exercise influences infiltration of Granzyme B+ cells in a transgenic mammary cancer model. **A.** Tumour initiation as age (day) of first indication of a palpable tumour in running (Runners) and non-running (Non-runners) MMTV-PyMT mice. n=11, ns = not significant, two-tailed t test. **B.** Left: tumour volume (mean and SEM) measured twice weekly for running PyMT mice and non-running PyMT mice. Right: survival curve. n=10-11, ns = not significant, Log-rank (Mantel-Cox) survival test. **C.** Tumour stage from histological scoring (1-4) in mammary glands of running and non-running PyMT mice at 12 (n=6-10) and 8 (n=4) weeks of age, ns = not significant, two-tailed t test. **D.** Individual body weight for WT and PyMT running and non-running mice. n=7-10, ns = not significant, one-way ANOVA with Tukey’s multiple comparison test. **E.** Immunohistological characterization of MMTV-PyMT tumours from non-running and running mice using CD3, F4/80, PCNA, Podocalyxin and granzyme B (GzmB) antibodies respectively. n=8-15, ns = not significant, * P <0.05, Two-tailed unpaired t test **F.** Frequency of myeloid populations within I3TC tumour. Ns = not significant, two-tailed t-test. **G.** Frequency of CD8+ T cells in spleens and I3TC tumours of animals treated with isotype control and anti-CD8 (*α*-CD8) depleting antibody.

**Supplementary Figure 2.**
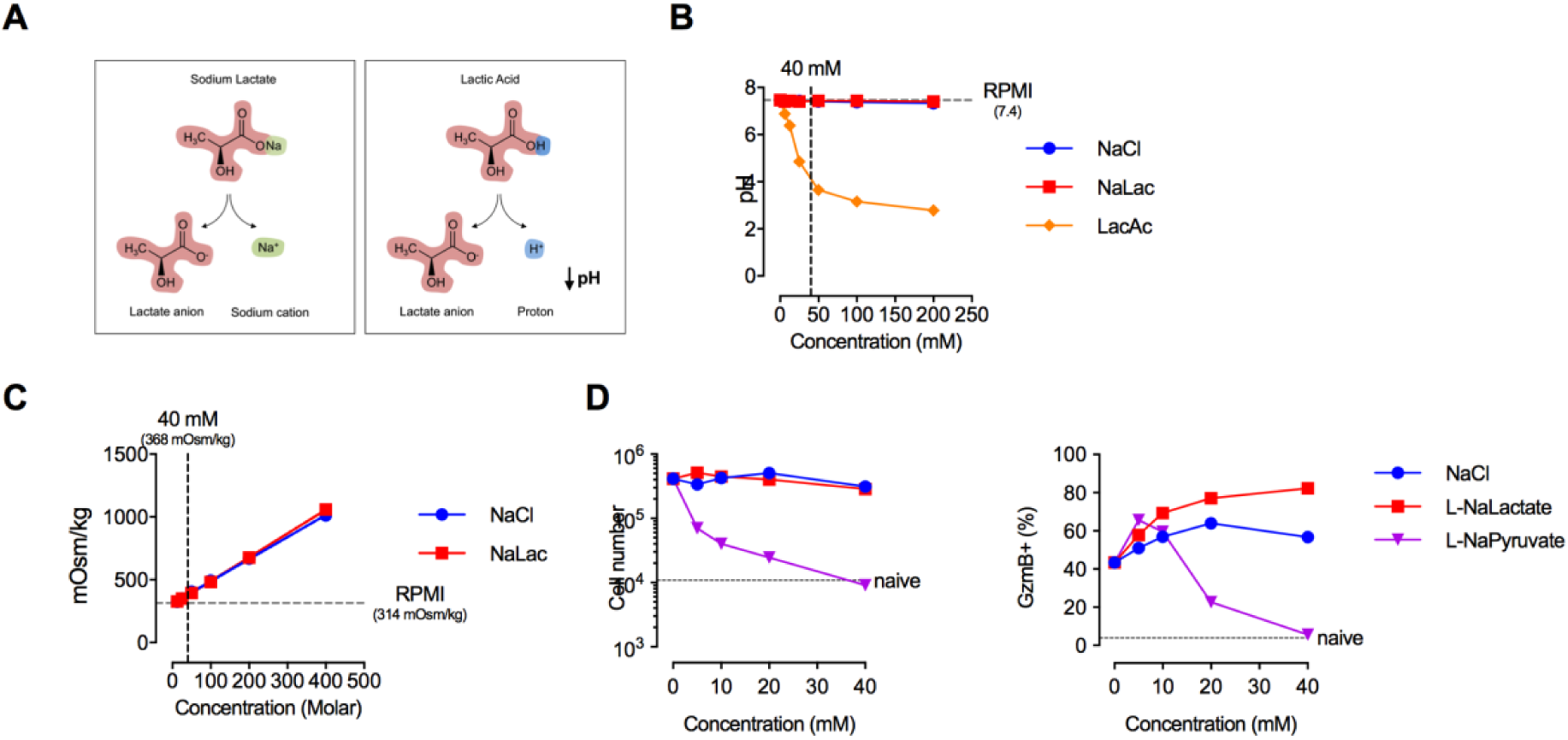
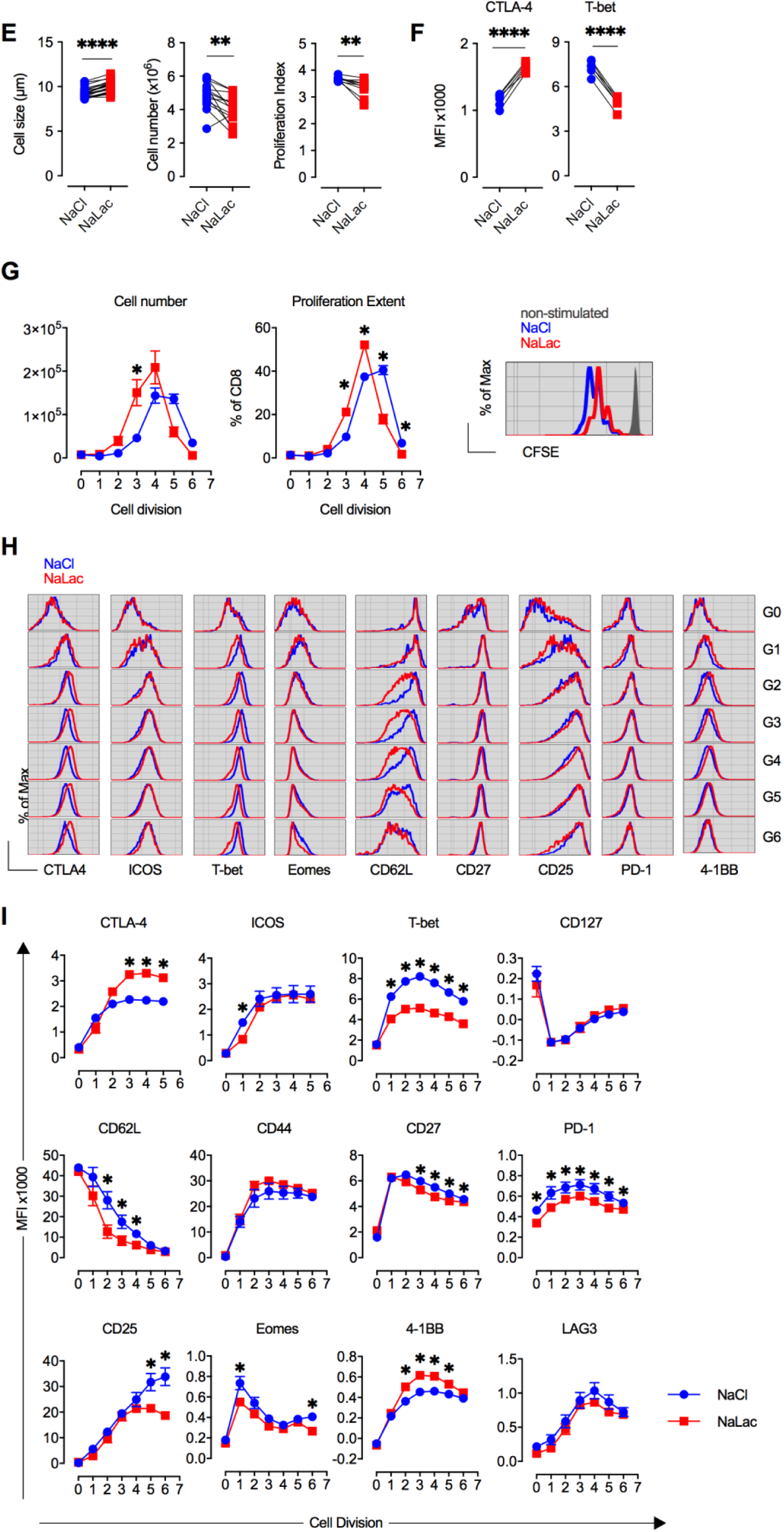
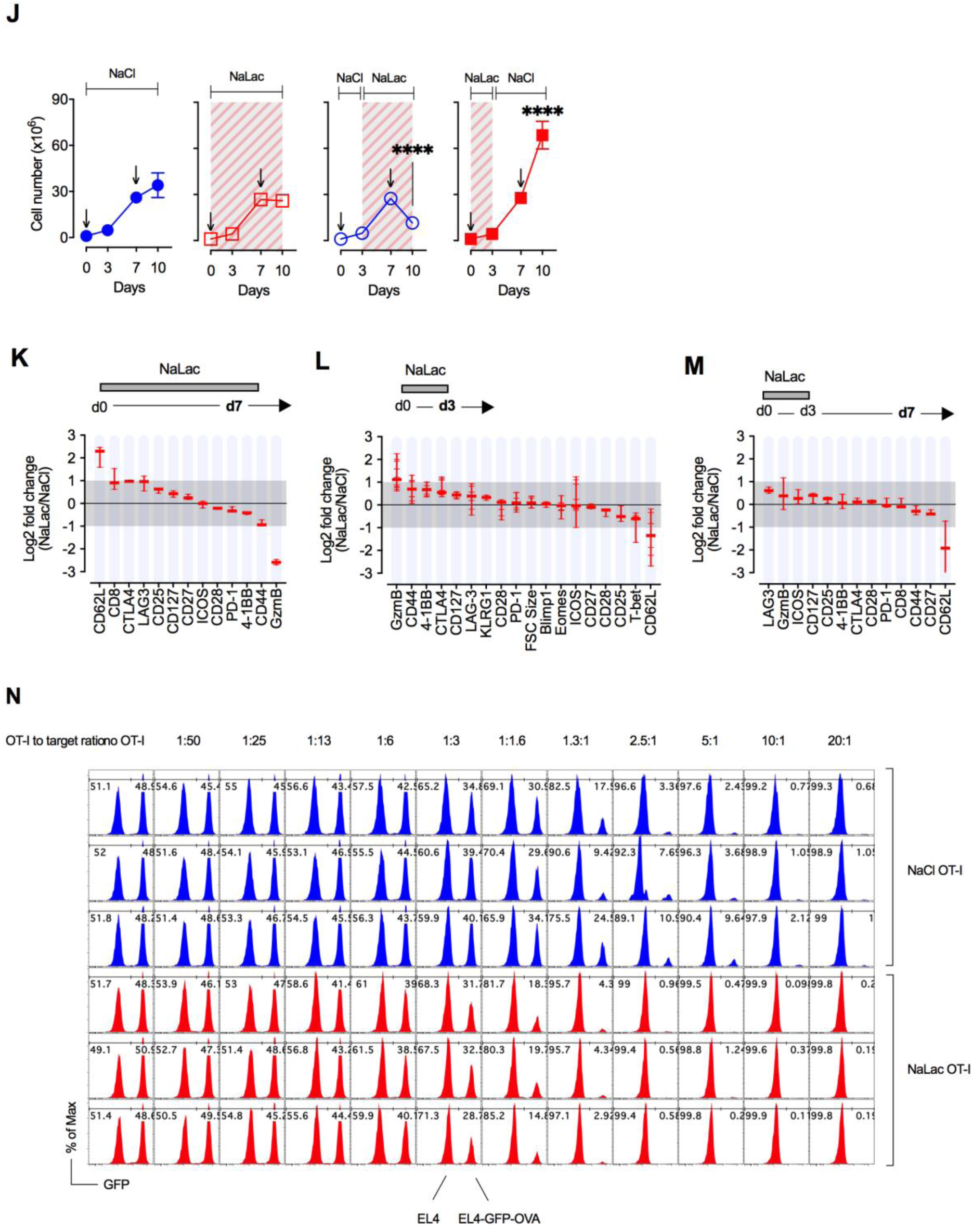
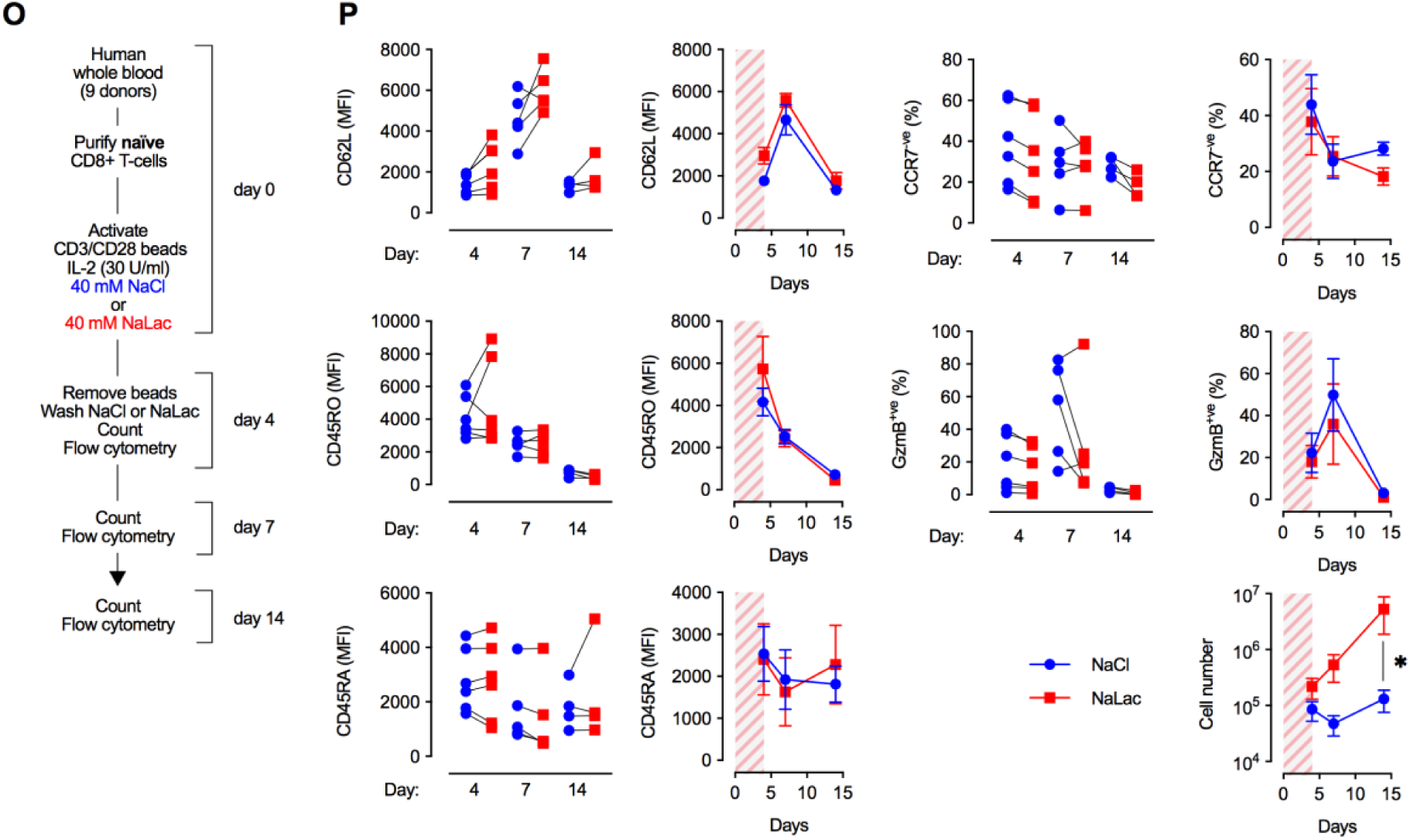
Lactate boosts proliferation and cytotoxic differentiation of CD8+ T cells. **A.** Diagram representing the dissociation of Sodium Lactate (NaLac, left) and Lactic Acid (LacAc, right) in aqueous solution. **B.** pH at room temperature of NaCl, NaLac and LacAc solutions in complete RPMI. Horizontal dotted line: RPMI reference. **C.** Osmolality of NaCl and Sodium Lactate (NaLac) solutions in complete RPMI. Horizontal dotted line: RPMI reference. Vertical dotted line: osmolality of 40 mM NaCl/NaLac solution in RPMI. **D.** Effect of varying concentrations of NaCl, L-NaLactate, and L-NaPyruvate on OT-I CD8+ T cell number and GzmB expression 72 hours after activation with SIINFEKL in the presence of IL-2. Dotted line: naive (non-activated). **E.** Mouse CD8+ T cells were activated with anti-CD3/CD28 microbeads and IL-2 for 72 hours in the presence of 40 mM Sodium Chloride (NaCl) or Sodium Lactate (NaLac). Effect of NaLac on cell size, cell number, and proliferation index,**F.** Effect of NaLac on intracellular expression of CTLA-4 and intranuclear expression of T-bet. For (A) and (B) each pair represents an independent mouse donor (n=6-21),**** P<0.0001, ** P<0.01, two-tailed paired t test. **G.** Absolute cell number (left) and frequency (middle) of CD8+ T cells at each cell generation as determined by CFSE dilution (right). Mean and SEM of n=3 independent mouse donors. * P<0.001 repeated-measures two-way ANOVA with Sidak’s multiple comparison test. **H.** Flow cytometry histograms showing fluorescence intensity of CTLA-4, ICOS, T-bet, Eomes, CD62L, CD27, CD25, PD-1 and 4-1BB at each cell division (G0-G6) 72 hours after activation with 40 mM NaCl (blue line) or 40 mM NaLac (red line). **I.** Median fluorescence intensity (MFI) at each cell generation summarizing data shown in (D). Mean and SEM of n=3-6 independent mouse donors. * P <0.001 repeated-measures two-way ANOVA with Sidak’s multiple comparison test. **J.** Cell number (mean and SEM) of mouse CD8+ T cells activated in the presence of interleukin-2 (IL-2) and different timings of exposure to 40 mM NaCl or NaLac (grey and red area). Cells were stimulated at days 0 and 7 (arrow). n=3 independent mouse donors, **** P <0.0001 repeated-measures two-way ANOVA with Dunnet’s multiple comparison test, comparing to no lactate treatment (filled blue circles). **K.** Ranked Log2 fold change of median fluorescence intensity of surface, cytoplasmic and nuclear CD8+ T cell differentiation proteins 7 days after activation in the presence of Sodium Lactate (NaLac) vs NaCl. Thick horizontal lines: median fold change. Thin horizontal lines: individual mouse donors. Vertical line: range. Grey area: range between 2-fold increase and 2-fold decrease. n=3 **L.** As described in K but analysis at day 3 after treatment with NaCl/NaLac. n=3-12. **M.** As described in K but analysis at day 7, 4 days after withdrawal of NaCl/NaLac. n=3 **N.** Mouse OVA-specific OT-I CD8+ T cells were activated with anti-CD3/CD28 microbeads and IL-2 for 72 hours in the presence of 40 mM Sodium Chloride (NaCl) or Sodium Lactate (NaLac). To measure cytotoxicity after activation OT-I cells were washed and counted before being co-cultured with a 1:1 mix of the thymoma cell line EL4 and a variant expressing OVA and GFP (EL4-GFP-OVA) for 24 hours. A constant number of EL4/EL4-GFP-OVA cells was co-cultured with an increasing number of OT-I cells (OT-I to EL4 ratio ranging from 1:50 to 20:1). Specific cytotoxicity was measured by flow cytometry by gating on EL4 cells (CD8-negative, SSC-and FSC-high) and measuring the frequency of the GFP-positive peak (OVA-positive, target peak) relative to the GFP-negative (OVA-negative, reference peak). EL4-GFP-OVA to EL4 ratio was normalised to that of cultures where no OT-I cells were added (0% cytotoxicity). Plots are flow cytometry histograms pre-gated on EL4 events and show GFP fluorescence peaks for 3 individual OT-I donors at varying OT-I to EL4 ratios. GFP high peak = EL4-GFP-OVA, GFP low peak = EL4. Numbers represent frequency of each peak. **O.** Experimental layout. Naïve CD8+ T cells were purified from whole blood of 9 healthy human donors and activated for 4 days with anti-CD3/CD28 beads, IL-2 and either 40 mM Sodium Chloride (NaCl) or 40 mM Sodium Lactate (NaLac). At day 4 activation beads were removed and cells washed and counted followed by phenotypic analysis by flow cytometry. Remaining CD8+ T cells were left in culture with IL-2 without further addition of NaCl or NaLac. Cell number and flow cytometry analysis were also performed at days 7 and 14. **P.** Cumulative cell number and median fluorescence intensity (MFI) of CD8+ T cell differentiation markers CD45RA, CD45RO and CD62L and frequency of CCR7-negative and GzmB-positive cells at days 4, 7 and 14. Left plots show paired data for individual donors at each time point (n=9 for cell number, n=4 for phenotypic markers). Right plots show mean and SEM over time for each parameter (grey/red shaded area = exposure to NaCl or NaLac). * P>0.05, two-tailed ratio paired t-test.

**Supplementary Figure 3.**
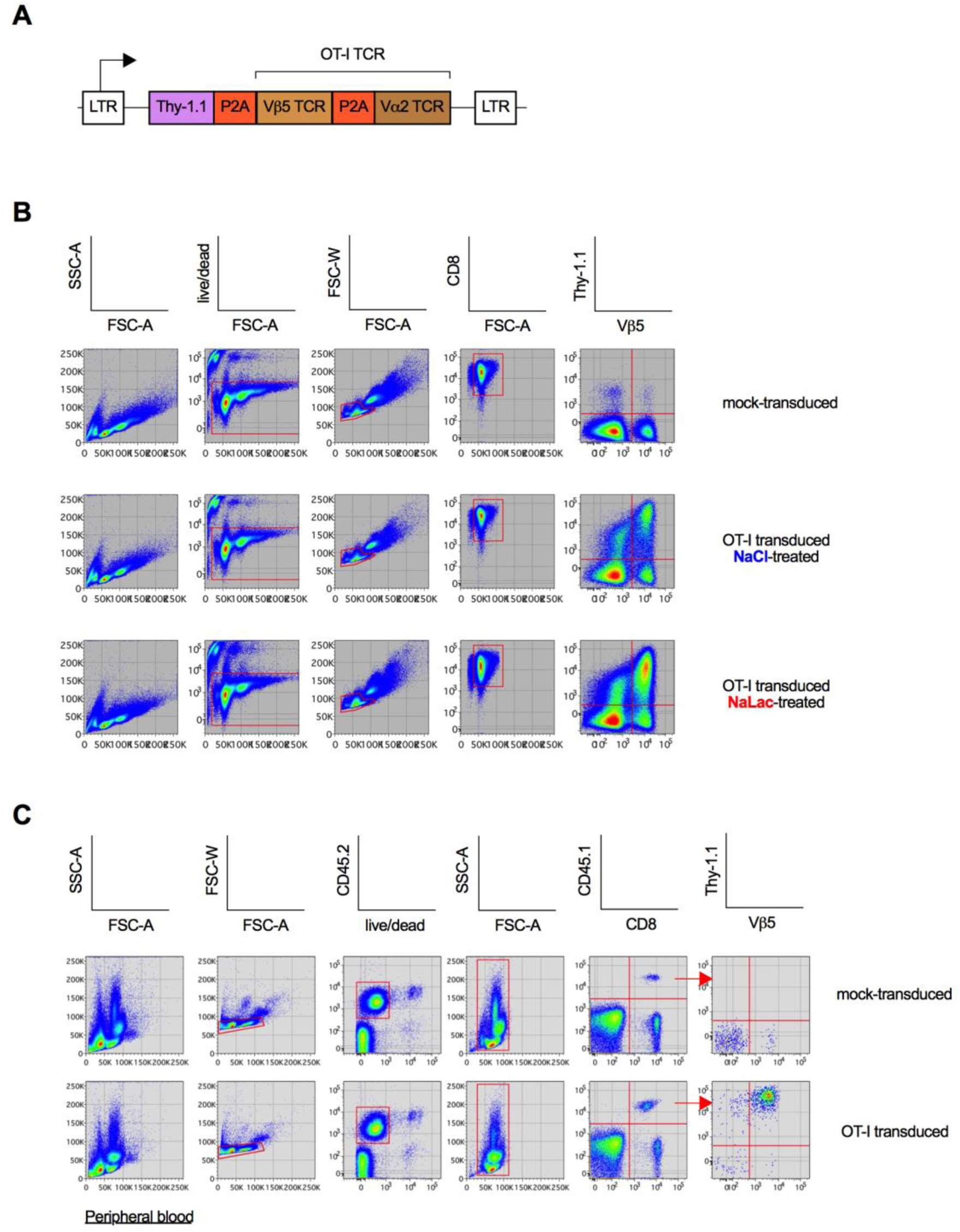
Sodium lactate enhances the *ex vivo* production of therapeutic CD8+ T cells. **A.** Layout of retroviral vector encoding a polycistronic peptide composed of the surface transduction marker Thy-1.1 and the TCR chains (V*α*2 and V*β*5) of the anti-OVA OT-I TCR. Elements are connected by self-cleaving picornavirus 2A sequences (P2A). LTR: retroviral long terminal repeats. **B.** Gating strategy to identify transduced CD8+ T cells in *ex vivo* culture. FSC-A, FSC-W: forward scatter area and width respectively; SSC-A: side scatter area; live/dead: near infra red live dead stain. **C.** Gating strategy to identify transduced CD8+ T cells in mouse peripheral blood 7 days after injection. CD45.2 and CD45.1: congenic markers for identification of endogenous and adoptively transferred immune cells, respectively.

**Supplementary Figure 4.**
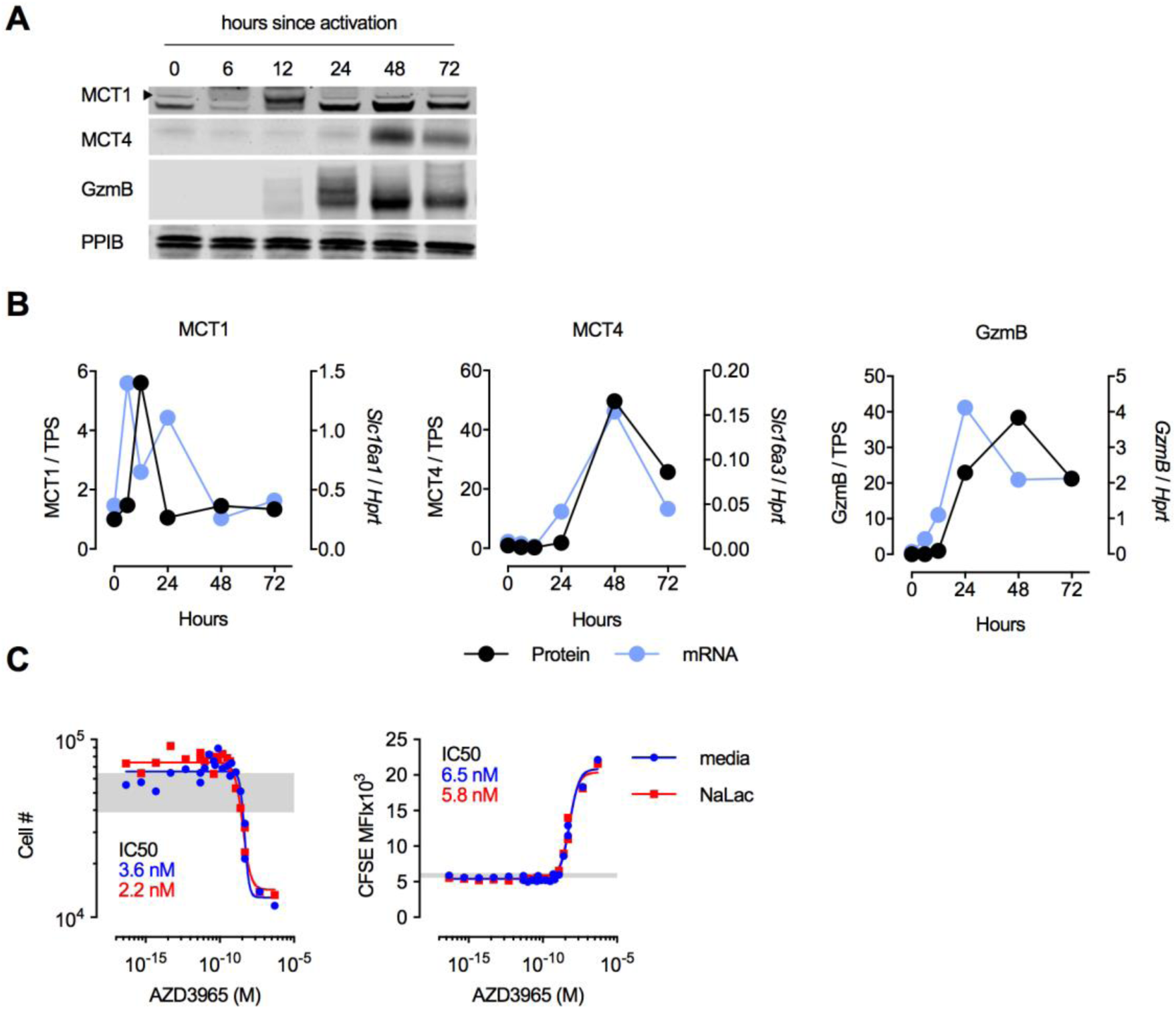
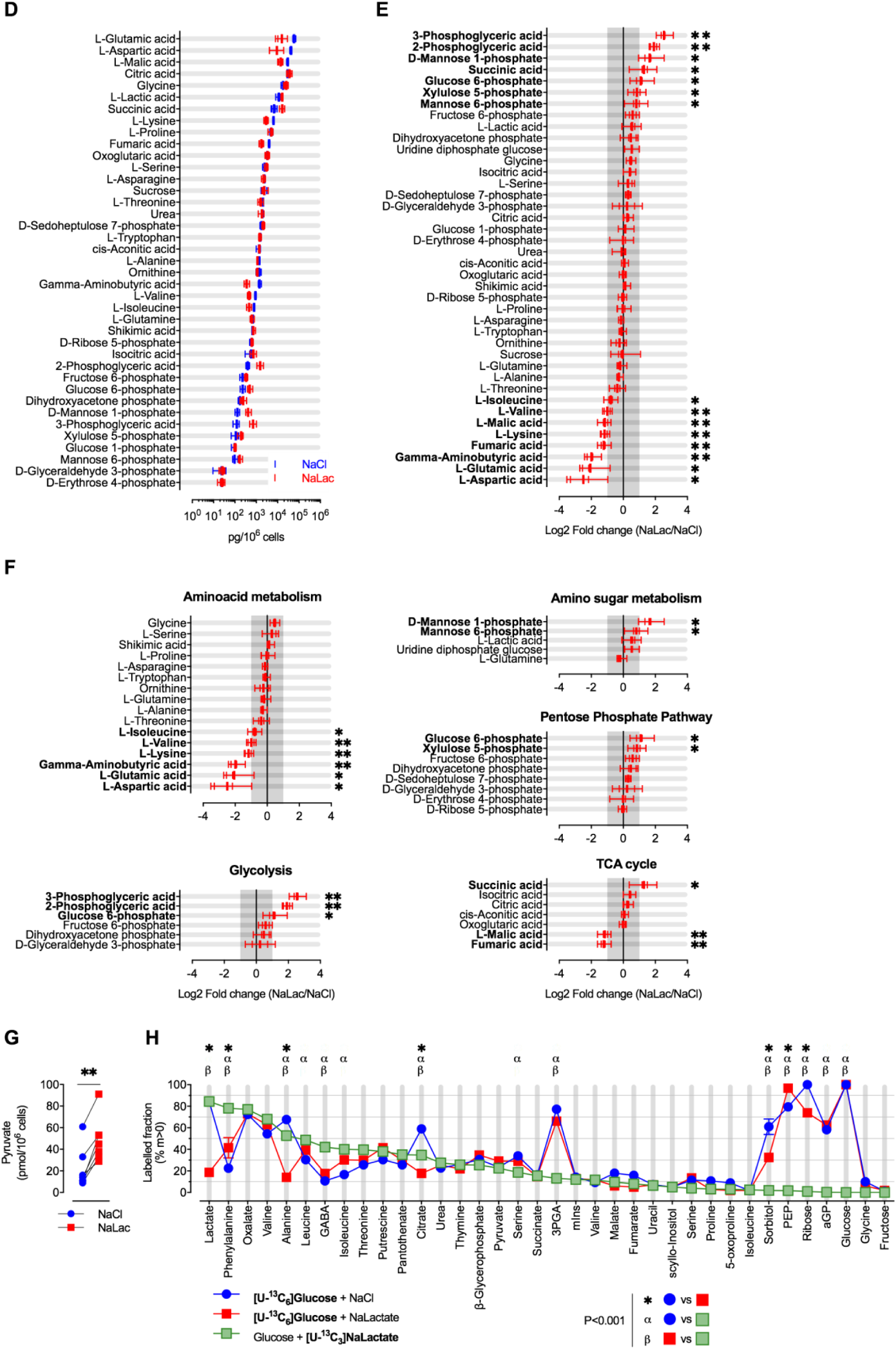
Lactate is utilized as a fuel by CD8+ T cells and alters the metabolic landscape. **A.** 15 µg of CD8+ T cell protein extracts from 0, 6, 12, 24, 48, and 72 hours post-activation were probed with antibodies against MCT1, MCT4, GzmB and PPIB (cyclophilin B). **B.** Protein (normalised to total protein stain; TPS) and mRNA (normalised to HPRT) levels of MCT1, MCT4 and GzmB over time since activation. **C.** CD8+ T cells were activated for 72 hours in the presence of the MCT1 inhibitor AZD3985. 40 mM Sodium Lactate (NaLac) or plain media were used. A non-linear fit ([Inhibitor] vs. response --Variable slope (four parameters)) was used to determine IC50. Grey area = range of untreated controls. **D.** Sugar Phosphate and GC-MS quantification of key metabolites in CD8+ T cells 72 hours after activation in the presence of 40 mM NaCl or Sodium Lactate (NaLac). Thick vertical lines: median amount of metabolite per 10^6^ cells. Thin vertical lines: individual mouse donors (n=4). Horizontal line: range. Metabolites are ranked from highest to lowest frequent in control (NaCl)-treated cells. **E.** Same dataset shown in A but expressed as Log2 fold change of metabolite quantity (NaLac vs NaCl). Thick vertical lines: median fold change. Thin vertical lines: individual mouse donors (n=4). Horizontal line: range. Grey area: range between 2-fold increase and 2-fold decrease.Metabolites are ranked from most upregulated to most downregulated in NaLac-treated cells. ** P<0.01, * P<0.05, two-tailed ratio paired t test. **F.** Same dataset and presentation as in B but grouping metabolites by pathway**G.** Pyruvate levels determined by a chemiluminescent assay. Each pair represents an independent mouse donor (n=7). ** P<0.01, two-tailed paired t-test. **H.** Mouse CD8+ T cells were activated for 72 hours in the presence of 11.1 mM [U-^13^C_6_]Glucose and 40 mM NaCl (blue circles), or 11.1 mM [U-^13^C_6_]Glucose and 40 mM NaLac (red squares), or 11.1 mM Glucose and 40 mM [U-^13^C_3_]NaLac (green squares). Proportion of metabolites in which at least one carbon is labelled (m>0) ranked from most to least labelled by [U-^13^C_3_]NaLac. n=4 independent mouse donors. *, *α, β* P<0.001, two-way ANOVA with Tukey’s multiple comparison test.

**Supplementary Figure 5.**
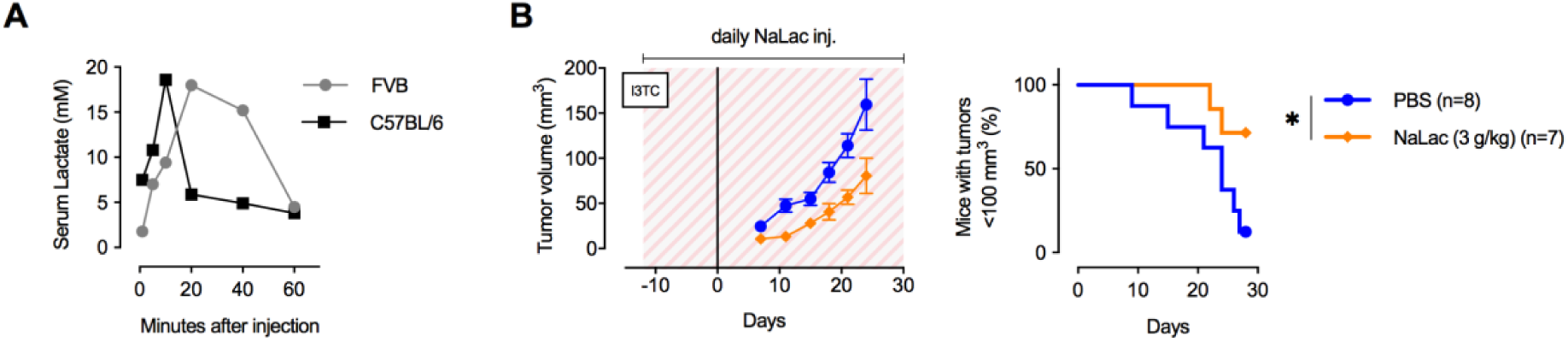
Daily administration of Sodium Lactate delays tumour growth. **A.** Peripheral blood lactate concentration (as measured with Accutrend® Plus) following a single dose of 2 g/kg Sodium Lactate (NaLac) administered i.p in FVB or C57BL/6J mice. **B.** Daily doses of PBS or 3 g/kg NaLac were administered i.p to FVB mice for 12 days before subcutaneous inoculation with with the breast cancer cell line I3TC. Injections were continued throughout the experiment. Graphs show tumour volume (mean and SEM) over time. * P <0.05, Log-rank (Mantel-Cox) survival test.

